# Structure of Zip2:Spo16, a conserved XPF:ERCC1-like complex critical for meiotic crossover formation

**DOI:** 10.1101/258194

**Authors:** Kanika Arora, Kevin D. Corbett

## Abstract

In eukaryotic meiosis, generation of haploid gametes depends on the formation of inter-homolog crossovers, which enable the pairing, physical linkage, and eventual segregation of homologs in the meiosis I division. A class of conserved meiosis-specific proteins, collectively termed ZMMs, are required for formation and spatial control of crossovers throughout eukaryotes. Here, we show that three *S. cerevisiae* ZMM proteins – Zip2, Zip4, and Spo16 – interact with one another and form a DNA-binding complex critical for crossover formation and control. We determined the crystal structure of a Zip2:Spo16 subcomplex, revealing a heterodimer structurally related to the XPF:ERCC1 endonuclease complex. Zip2:Spo16 lacks an endonuclease active site, but binds specific DNA structures found in early meiotic recombination intermediates. Mutations in multiple DNA-binding surfaces on the Zip2:Spo16 complex severely compromise DNA binding, supporting a model in which the complex’s central and HhH domains cooperate to bind DNA. Overall, our data support a model in which the Zip2:Zip4:Spo16 complex binds and stabilizes early meiotic recombination intermediates, then coordinates additional factors to promote crossover formation and license downstream events including synaptonemal complex assembly.

## Introduction

Sexual reproduction in eukaryotes requires the production of haploid gametes in a specialized cell-division program called meiosis, followed by the fusion of two gametes to generate diploid offspring. The reduction of ploidy in meiosis is enabled by the formation of crossovers or chiasmata, specific inter-homolog recombination events that physically link homologs and enable their segregation in the meiosis I division. Because of their importance for accurate chromosome segregation, the formation of crossovers (COs) is subject to tight spatial and temporal regulation: overlapping feedback pathways ensure that each homolog pair receives at least one CO, and that the overall number of COs is kept within a tight range. Most COs are also subject to “interference”; that is, they are spaced farther apart along chromosomes than expected by random chance, implying a regulatory mechanism that communicates the status of recombination along chromosomes (Wang *et al*, 2015; Berchowitz & Copenhaver, 2010). Defects in the production or spatial regulation of COs can lead to chromosome mis-segregation and aneuploidy, a major cause of miscarriage and developmental disorders in humans (Hassold *et al*, 2007; Hassold & Hunt, 2001).

While the molecular machinery that mediates meiotic recombination is highly conserved and well-understood, the regulatory mechanisms controlling the number and spatial distribution of COs are less well-characterized. In the budding yeast *S. cerevisiae*, entry into meiotic prophase is accompanied by assembly of the meiotic chromosome axis, which organizes each pair of replicated sister chromosomes as a linear array of chromatin loops and promotes the formation of DNA double-strand breaks (DSBs) along each chromosome by the Spo11 endonuclease (Mao-Draayer *et al*, 1996; Xu *et al*, 1997; Peciña *et al*, 2002; Schwacha & Kleckner, 1997; Woltering *et al*, 2000; Blat *et al*, 2002; Keeney *et al*, 1997). These DSBs are resected to free 3’ single-stranded ends and loaded with two related recombinases, Rad51 and Dmc1, which mediate the invasion of a homologous DNA duplex to form a single-end invasion or D-loop intermediate (Bishop *et al*, 1992; Cloud *et al*, 2012). In meiotic prophase, use of the identical sister chromatid as a repair template is strongly inhibited by the chromosome axis, thereby promoting invasion of the homolog instead (Niu *et al*, 2005; Subramanian *et al*, 2016; Carballo *et al*, 2008; Niu *et al*, 2009; 2007; Schwacha & Kleckner, 1997; Lao *et al*, 2013). After initial invasion of the homolog and formation of a D-loop, several competing pathways vie to determine the fate of this early recombination intermediate. Often, the D-loop is dissolved by the combined action of DNA topoisomerases and helicases (in *S. cerevisiae*, the Sgs1, Top3, and Rmi1 proteins)(Kaur *et al*, 2015; Oh *et al*, 2007; De Muyt *et al*, 2012). If the free 3’ end has undergone new DNA synthesis past the original break, it may re-anneal with the other broken end and be repaired to give a non-crossover (NCO); this pathway is referred to as synthesis-dependent strand annealing (SDSA) and is responsible for the bulk of NCO formation (Andersen & Sekelsky, 2010; Allers & Lichten, 2001). A subset of strand invasion intermediates are further processed into double Holliday Junction (dHJ) intermediates (Allers & Lichten, 2001; Hunter & Kleckner, 2001; Schwacha & Kleckner, 1995), which may become either COs or NCOs. Type 1 “interfering” COs are generated by specific cleavage of dHJs by the Mlh1:Mlh3 endonuclease (Argueso *et al*, 2004; Zakharyevich *et al*, 2012). A minor competing pathway involving non-specific endonucleases (in *S. cerevisiae*, Mus81:Mms4, Yen1, and Slx1:Slx4) can generate either NCOs or type 2 “noninterfering” COs (Argueso *et al*, 2004; Blanco *et al*, 2014; de los Santos *et al*, 2003; Oh *et al*, 2008; Zakharyevich *et al*, 2012; De Muyt *et al*, 2012).

The formation of type 1 COs is tightly regulated by a group of proteins collectively termed “ZMM” proteins after their gene names in *S. cerevisiae*: Zips (Zip1 / Zip2 / Zip3 / Zip4), Msh4:Msh5, and Mer3 (Lynn *et al*, 2007). Most ZMM proteins are conserved throughout eukaryotes, and their roles in promoting CO formation are generally well-outlined: for example, Mer3 is a DNA helicase (Mazina *et al*, 2004) and Msh4:Msh5, relatives of the MutS family DNA mismatch recognition proteins, are proposed to bind and stabilize a branched recombination intermediate and recruit the Mlh1:Mlh3 endonuclease for specific dHJ cleavage (Snowden *et al*, 2008; 2004). Zip3 is related to ubiquitin / SUMO E3 ligase proteins (Perry *et al*, 2005), and as such likely has a role in regulation of protein complex formation or degradation at crossover sites (Ahuja *et al*, 2017; Rao *et al*, 2017). Zip1 is the major component of “transverse filaments” within the synaptonemal complex, a conserved structure that nucleates at initial homolog interaction sites including centromeres and recombination sites, then extends to bring each homolog pair into close juxtaposition along their lengths (Sym *et al*, 1993; Dong & Roeder, 2000). The synaptonemal complex is important for the resolution of crossovers (Zickler & Kleckner, 1999; Sym *et al*, 1993; Dong & Roeder, 2000) and its assembly is also coordinated with removal of Hop1 from the axis by the AAA + ATPase Pch2 (Chen *et al*, 2014; Joshi *et al*, 2009; Borner *et al*, 2008; Ye *et al*, 2017), an important feedback mechanism that limits further DSB and CO formation.

Of the identified ZMM proteins in *S. cerevisiae*, the roles of Zip2, Zip4, and their more recently-identified binding partner Spo16 are the least well-understood. These three proteins localize to recombination sites on meiotic chromosomes (Fung *et al*, 2004; Chua & Roeder, 1998) and depend on one another for chromosome localization (Tsubouchi *et al*, 2006; Shinohara *et al*, 2008), suggesting that they act together as a complex. All three proteins are required for wild-type levels of COs (Chua & Roeder, 1998; Shinohara *et al*, 2008; Malavasic & Elder, 1990; Tsubouchi *et al*, 2006), *𝓏ip2* and *𝓏ip4* mutants show defects in crossover interference (Tsubouchi *et al*, 2006; Chen *et al*, 2008), and *spo16* and *𝓏ip2* mutants also show defects in the formation of single-end invasion / D-loop and dHJ recombination intermediates (Shinohara *et al*, 2008; Börner *et al*, 2004). Together, these data suggest that Zip2, Zip4, and Spo16 may directly promote the formation of type 1 COs, potentially by aiding the formation of early recombination intermediates or stabilizing these intermediates against disassembly. Similar findings that both the *A. thaliana* and mammalian Zip2 homologs (SHOC1/SHOC1) are required for wild-type levels of crossovers in these organisms (Macaisne *et al*, 2008; Guiraldelli *et al*, 2018) suggest that a Zip2-containing complex is a highly-conserved and critical player in meiotic CO formation throughout eukaryotes.

In addition to their effects on meiotic recombination, Zip2, Zip4, and Spo16 also play a key role in synaptonemal complex assembly: in mutants of all three genes, the synaptonemal complex protein Zip1 forms foci at recombination sites, but does not then extend along chromosomes to form the full synaptonemal complex (Chua & Roeder, 1998; Tsubouchi *et al*, 2006; Shinohara *et al*, 2008). Zip2 and Zip4 have also been observed to localize at the ends of Zip1 stretches, both on synapsed chromosomes and on extra-chromosomal Zip1 assemblies termed polycomplexes (Tsubouchi *et al*, 2006; Chua & Roeder, 1998). Recently, mass spectrometry of Zip2:Zip4:Spo16 complexes purified from meiotic cells has indicated that these proteins directly or indirectly interact with the chromosome axis proteins Hop1 and Red1, the axis remodeler Pch2, and the Msh4:Msh5 complex (De Muyt *et al*, 2018). Overall, these findings suggest that Zip2, Zip4, and Spo16 interact with other ZMMs as well as both chromosome axis and synaptonemal complex proteins, and thereby play a central role in coordinating CO formation with chromosome axis remodeling and synaptonemal complex assembly.

*S. cerevisiae* Zip2 and its homologs in *Arabidopsis thaliana* (AI481877/SHOC1/ZIP2H) and mammals (C9ORF84/SHOC1) show weak homology to XPF, a structure-specific endonuclease that plays important roles in nucleotide excision repair with its binding partner, ERCC1 (Macaisne *et al*, 2011; 2008; De Muyt *et al*, 2018). More recently, De Muyt et al. found that Spo16 binds the XPF-like domain of Zip2 specifically and shows weak homology to ERCC1, suggesting that these proteins may form a XPF:ERCC1-like complex (De Muyt *et al*, 2018). Here, we outline the architecture of the *S. cerevisiae* Zip2:Zip4:Spo16 complex and show by x-ray crystallography that Zip2 and Spo16 indeed form an XPF:ERCC1-like heterodimer. Zip2:Spo16 lacks an endonuclease active site, suggesting that it instead binds specific DNA structures and nucleates assembly of a larger regulatory complex, similar to the related FANCM:FAAP24 DNA repair complex (Yang *et al*, 2013). We show that, like XPF:ERCC1, Zip2:Spo16 can bind a variety of DNA structures, including single – strand/double-strand DNA junctions with the geometry found in meiotic strand-invasion intermediates, and branched/bent DNA structures. We propose a model in which the Zip2:Zip4:Spo16 complex cooperates with Msh4:Msh5 to recognize and stabilize early recombination intermediates, recruit downstream crossover factors to promote the formation of type 1 COs, and coordinate the formation of COs with assembly of the synaptonemal complex.

## Results

### Architecture of the Zip2:Zip4:Spo16 complex

Zip2, Zip4, and Spo16 share similar roles in promoting meiotic crossover formation and synaptonemal complex assembly, co-localize on meiotic chromosomes, and depend on one another for their chromosome localization (Tsubouchi *et al*, 2006; Shinohara *et al*, 2008). Recently, De Muyt et al. showed that the three proteins can be co-purified from meiotic cell lysate, directly interact in yeast two-hybrid assays, and depend on one another for localization to DNA double-strand breaks sites (De Muyt *et al*, 2018). To complement this work and comprehensively outline interactions between Zip2, Spo16, and Zip4, we first used yeast two-hybrid analysis. We found that Spo16 binds the conserved C-terminal region of Zip2 (residues 499-704), which contains Zip2’s predicted XPF-like domain (**Figure 1A,B**). We also found that Zip2’s less well-conserved N-terminal region (residues 1-499) interacts with full-length Zip4, though we detected this interaction with only one tag configuration (**Figure 1B**). Given that the Zip2 C-terminal region interacts with Spo16 and may require this interaction for solubility, we next performed yeast three-hybrid assays with one vector encoding both Zip2 and Spo16, and a second vector encoding Zip4. We detected a strong interaction in this assay that was dependent on the N-terminal region of Zip2 (**Figure 1C**). By progressively truncating the Zip2 N-terminus, we found that removal of the N-terminal 200 residues of Zip2 does not affect Zip4 binding, while removal of 287 residues partially disrupts binding, and removal of 380 residues completely disrupts binding. While the N-terminal region of Zip2 is mostly poorly conserved, this region does contain several short, highly-conserved motifs (**Figure S1**). Based on our yeast three-hybrid data, two conserved motifs spanning residues 268-282 and 301-317 may be directly involved in Zip4 binding. Together, these data strongly indicate that Zip2 is the key central component of the Zip2:Zip4:Spo16 complex, with its N-terminal domain binding Zip4 and its C-terminal domain binding Spo16. Efforts to identify the Zip2-binding region of Zip4 by deletion analysis were unsuccessful, likely due to structural disruption when truncating the protein’s predicted array of TPR repeats (not shown).

**Figure 1.**
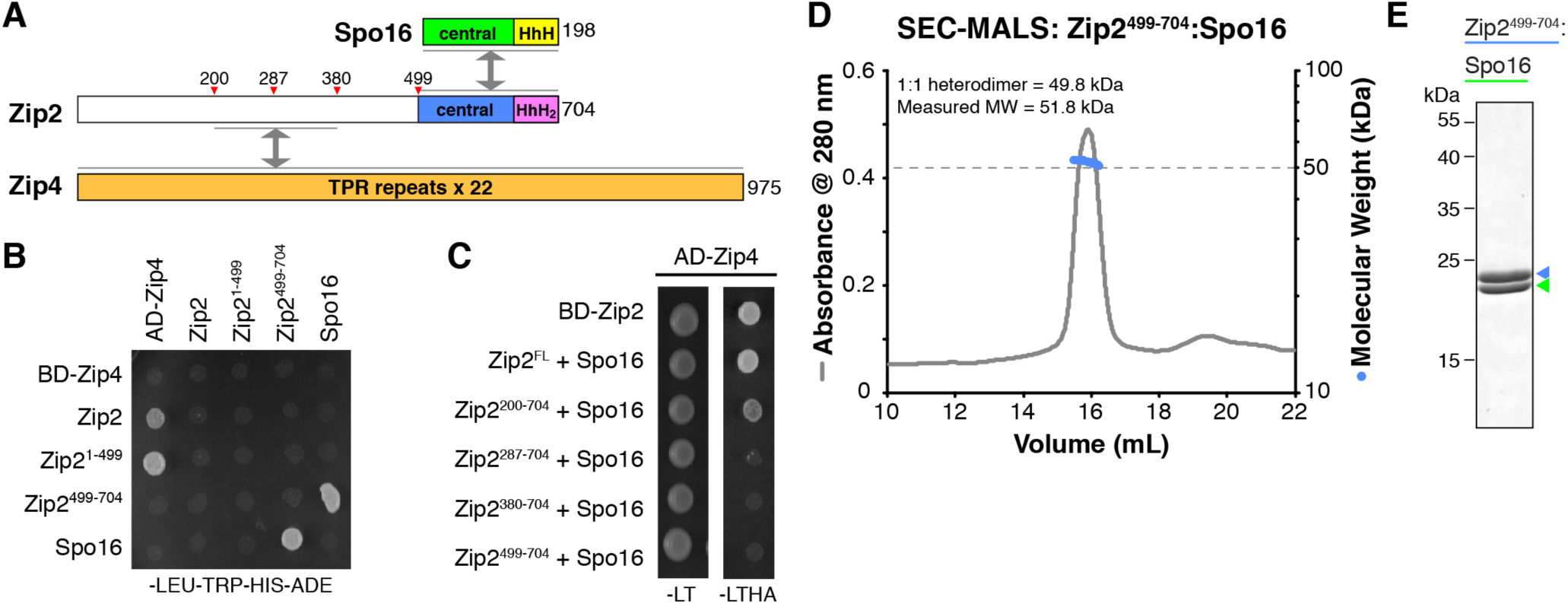
Zip2, Zip4, and Spo16 form a complex. (A) Domain structure of Zip2, Zip4, and Spo16. Gray arrows indicate interactions identified by yeast two-hybrid and yeast three-hybrid analysis. The N-terminal domain of Zip2 has been proposed to contain a WD40 *β*-propeller domain (Perry *et al*, 2005), but the region is poorly conserved (**Figure S1**) and modern structure-prediction algorithms do not support this assignment. HhH: helix-hairpin-helix, HhH_2_: tandem helix-hairpin-helix. (B) Yeast two-hybrid analysis of interactions between Zip2, Zip4, and Spo16. See **Figure S2A** for complete results. (C) Yeast three-hybrid analysis. See **Figure S2B** for complete results. (D) Size exclusion chromatography/multi-angle light scattering (SEC-MALS) analysis of purified Zip2^499-704^:Spo16. The measured molecular weight (51.8 kDa) is consistent with a 1:1 heterodimer (molecular weight 49.8 kDa). (E) SDS-PAGE analysis of purified Zip2^499-704^:Spo16.

### The Zip2:Spo16 structure reveals an XPF:ERCC1-like complex

We next co-expressed Zip2 and Spo16 in *E. coli* for structural and biochemical analysis. While full-length Zip2 expressed poorly and was mostly insoluble, even when co-expressed with Spo16, we could co-express and purify the Zip2 XPF-like region (residues 499-704) with Spo16. The Zip2^499-704^:Spo16 complex forms a well-behaved 1:1 heterodimer (**Figure 1D,E**). While initial crystallization efforts were unsuccessful, we obtained crystals of the complex after methylation of surface lysine residues by formaldehyde/dimethylamine borane complex treatment (**Figure S3A**) (Walter *et al*, 2006; Kim *et al*, 2008). The resulting crystals (termed Form 1 hereon) adopted space group C2 and contained four copies of the complex per asymmetric unit. Extensive efforts to determine the structure of Form 1 crystals using heavy-atom derivatives failed due to translational pseudo-symmetry in these crystals.

To identify additional crystal forms for the Zip2^499-704^:Spo16 complex, we turned to surface entropy reduction, a strategy to promote crystallization by mutating short stretches of large polar residues to alanine (Derewenda, 2004). We mutated the highest-scoring segment in Zip2 identified by the UCLA SERp Server (Goldschmidt *et al*, 2007), residues 641-643, from the sequence KEK to AAA (**Figure S1**). The mutated complex, Zip2^499-704^SER:Spo16, behaved equivalently to wild-type protein in vitro but crystallized in several new conditions without surface lysine methylation. We optimized one condition in space group P2_1_2_1_2_1_ (termed Form 2 hereon), which contained one copy of the Zip2^499-704^SER:Spo16 complex per asymmetric unit, and determined the structure by single-wavelength anomalous diffraction methods (**Figure 2A, Table S1**). We then used molecular replacement to determine the Form 1 structure, yielding a total of five crystallographically-independent views of the Zip2^499-704^:Spo16 complex. These five structures are nearly identical (overall C*α* r.m.s.d. 0.3-1.4 Å; **Figure S3F**), indicating that neither crystallization strategy – surface lysine methylation or surface entropy reduction – significantly alters the complex’s structure.

**Figure 2.**
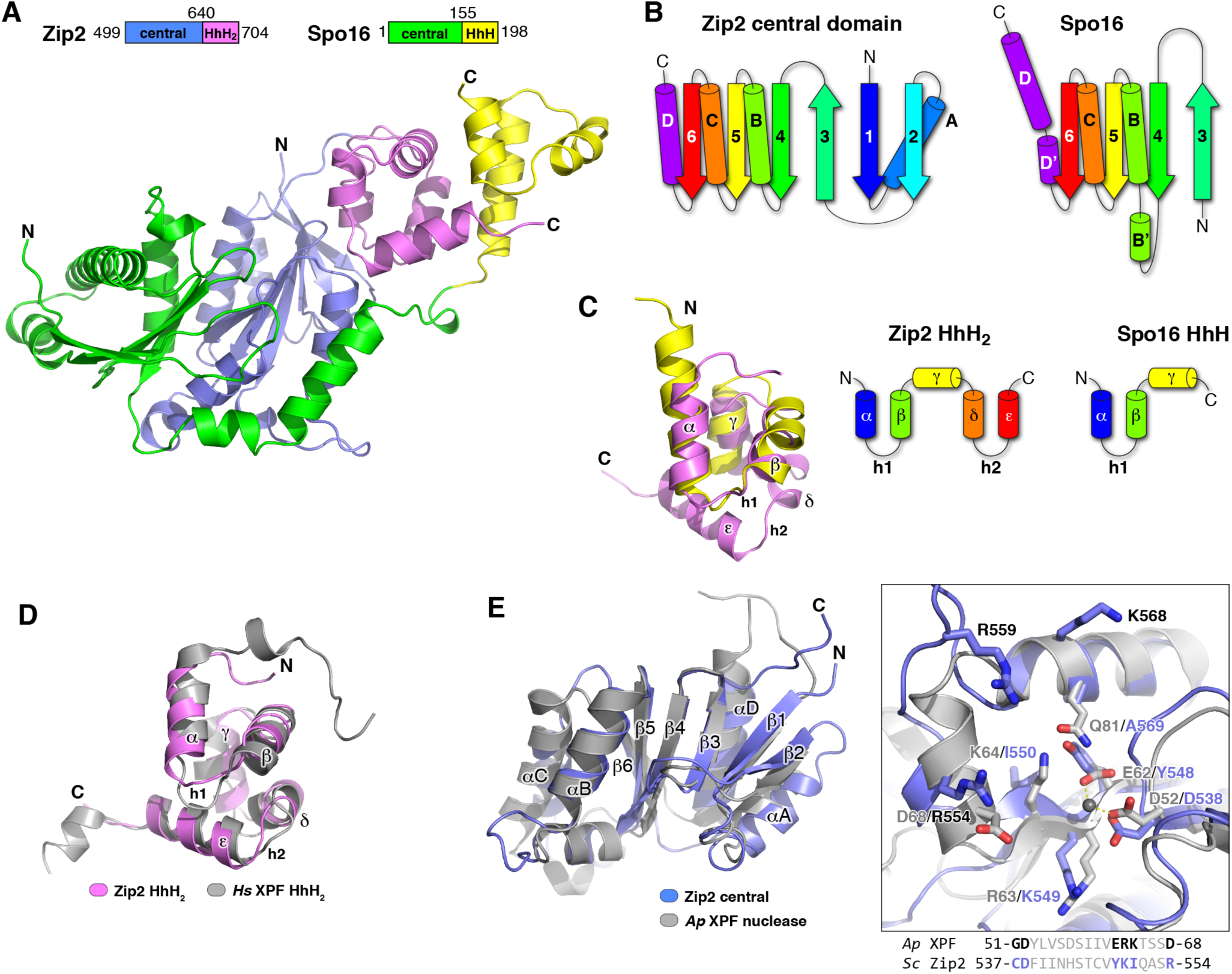
Structure of the Zip2^499-704^:Spo16 complex. (A) Domain schematic of Zip2 and Spo16 (top) and overall structure of the Zip2^499-704^:Spo16 dimer. See **Figure S3F** for an overlay of the five crystallographically-independent views of the dimer. (B) Secondary structure of the Zip2 and Spo16 central domains. *(C) Left:* Overlay of the Zip2 HhH_2_ and Spo16 HhH domains. C*α* r.m.s.d. = 2.65 Å over 34 atom pairs. HhH #1 comprises helices *α* and *β* separated by hairpin 1 (h1), and HhH #2 (not shared by Spo16) comprises helices *δ* and *ε*, separated by hairpin 2(h2). *Right:* secondary structure of the Zip2 HhH_2_ and Spo16 HhH domains. (D) Overlay of the Zip2 HhH_2_ domain (pink) with the HhH_2_ domain of *H. sapiens* XPF (PDB ID 1Z00; gray). C*α* r.m.s.d. = 1.97 Å over 51 atom pairs. (E) Overlay of the Zip2 central domain (blue) with the *Aeropyrum pernix* XPF nuclease domain (gray) (PDB ID 2BGW) (Newman *et al*, 2005). C*α* r.m.s.d. = 2.50 Å over 107 atom pairs. *Left:* close-up of the active site of *Ap* XPF (gray, with bound Mg^2+^ ion shown as a sphere) with the equivalent region of the Zip2 central domain (blue). Zip2’s DNA binding patch 1 (see **Figure 4B**) comprises residues R554, R559, and K568 (labeled in black).

The overall structure of Zip2^499-704^:Spo16 is similar to known XPF:ERCC1 and related complexes (**Figure 2A**). Zip2 possesses an XPF-like central domain (residues 499-640) and a C-terminal tandem helix-hairpin-helix (HhH_2_) domain (residues 641-704). Spo16 shows a similar two-domain structure, with an N-terminal central domain similar in fold to ERCC1, though significantly diverged (**Figure 2B**). As previously predicted by De Muyt et al. (De Muyt *et al*, 2018), the Spo16 C-terminal domain contains only a single HhH motif as opposed to the tandem HhH motifs found in ERCC1 (**Figure 2C,D**). The Zip2 and Spo16 central domains form a tight pseudo-symmetric dimer, with flexible linkers connecting these domains to the C-terminal HhH domains, which also form a tight dimer. The conformations of the linker regions and the relative positions of the central and HhH domains are identical in the five independent views of the Zip2^499-704^:Spo16 complex (**Figure S3F**), suggesting that the complex is relatively rigid, at least in the absence of DNA (see below). The juxtaposition of central and HhH domains is also similar to a recent structure of the human FANCM:FAAP24 complex, an inactive XPF:ERCC1-like complex with a key role in scaffolding the Fanconi Anemia “core” DNA-repair complex (Yang *et al*, 2013).

The key biochemical activities of XPF:ERCC1 are structure-specific DNA binding and single-strand DNA cleavage, primarily at single-strand—double-strand DNA junctions, as part of its role in the nucleotide excision repair pathway (Ciccia *et al*, 2008). DNA cleavage is catalyzed at a highly-conserved active site on the XPF central/nuclease domain. Prior sequence analyses of Zip2 and both *A. thaliana* and mammalian SHOC1 have suggested that these proteins likely lack endonuclease activity (Macaisne *et al*, 2008; De Muyt *et al*, 2018; Guiraldelli *et al*, 2018), and Zip2 was shown to lack endonuclease activity in vitro on a typical XPF:ERCC1 substrate (De Muyt *et al*, 2018). We overlaid the Zip2 central domain with the structure of *Aeropyrum pernix* XPF nuclease domain (Newman *et al*, 2005), and examined the region around the putative active site (**Figure 2E**). XPF-family proteins possess a highly-conserved motif, GDx_n_ERKx_3_D, which contains residues responsible for Mg^2^+ ion binding and DNA cleavage (Nishino *et al*, 2003). Of the highly-conserved residues in the XPF active-site motif, Zip2 possesses only one, D538, which corresponds to the first aspartate in the conserved XPF motif (D52 in *A. pernix* XPF; **Figure 2E, S1**). The region of Zip2 corresponding to the ERKx_3_D motif of XPF proteins does not contain any of these residues, and is highly variable across fungal Zip2 orthologs (**Figure 2E, S1**). Further, when we determined a structure from crystals grown in the presence of 10 mM MgCl_2_, we saw no evidence of a bound Mg^2+^ ion in this site (not shown). Thus, despite the conservation of one key active site residue with XPF, our structural evidence overall points to Zip2 lacking endonuclease activity, in agreement with prior biochemical data (De Muyt *et al*, 2018). This finding is again reminiscent of the FANCM:FAAP24 complex, in which the XPF ortholog FANCM also lacks endonuclease activity. Since FANCM:FAAP24 is thought to bind specific DNA structures and scaffold the assembly of a multi-subunit DNA repair complex (Yang *et al*, 2013), a similar DNA binding/scaffolding activity could explain existing genetic data on the roles of *ZIP2* and *SPO16* in CO formation.

### Zip2:Spo16 binds a range of DNA structures

The structural similarity of Zip2^499-704^:Spo16 to XPF:ERCC1 and related complexes including FANCM:FAAP24 and Mus81:Eme1 strongly suggests a structure-specific DNA binding function for this complex. Further, the delay in formation of single-end invasion and double Holliday Junction intermediates in *spo16* mutants (Shinohara *et al*, 2008) suggests that the complex may bind and stabilize a specific recombination intermediate to promote crossover formation. To test for DNA binding activity, we performed quantitative electrophoretic mobility-shift assays (EMSA) with reconstituted Zip2^499-704^:Spo16 and DNA substrates containing structural elements found in meiotic recombination intermediates. We first tested binding of Zip2^499-704^:Spo16 to a 40-base single-stranded DNA (ssDNA) and a 40-bp double-stranded DNA duplex (dsDNA). Zip2^499-704^:Spo16 bound robustly to dsDNA (*K*_*d*_ = 14 μM; **Figure 3A**) but barely interacted with ssDNA (**Figure S4A**). We next tested binding to ssDNA-dsDNA junctions, as XPF:ERCC1 family complexes recognize these structures for cleavage, and because strand invasion during meiotic homologous recombination generates a D-loop intermediate that features a ssDNA-dsDNA junction with a 5’ overhang. We found that Zip2^499-704^:Spo16 binds more strongly to a 5’-overhang junction than to either dsDNA or a junction with a 3’ overhang (5’ overhang *K*_*d*_ = 9 μM, 3’ overhang *K*_*d*_ = 17 μM; **Figure 3B, C**). Curiously, Zip2^499-704^:Spo16 bound more tightly still to a nicked DNA substrate (*K*_*d*_ = 6 μM; **Figure 3D**), suggesting that the complex may preferentially bind highly bent or bendable DNA. Finally, we tested binding of a Holliday Junction (HJ) substrate with 20-bp arms. While Zip2:Zip4:Spo16 localizes to meiotic recombination sites prior to the formation of the double Holliday Junction intermediate, binding to this substrate could indicate specificity for other DNA structures formed prior to the double Holliday Junction, or a general affinity for branched or bent structures. We observed robust binding of Zip2^499-704^:Spo16 to the HJ substrate, comparable to the nicked substrate (*K*_*d*_= 5 μM; **Figure 3E**). We also observed multiple shifted bands on our gels with the HJ substrate, indicating that multiple Zip2^499-704^:Spo16 complexes may bind a single HJ at high protein concentrations.

**Figure 3.**
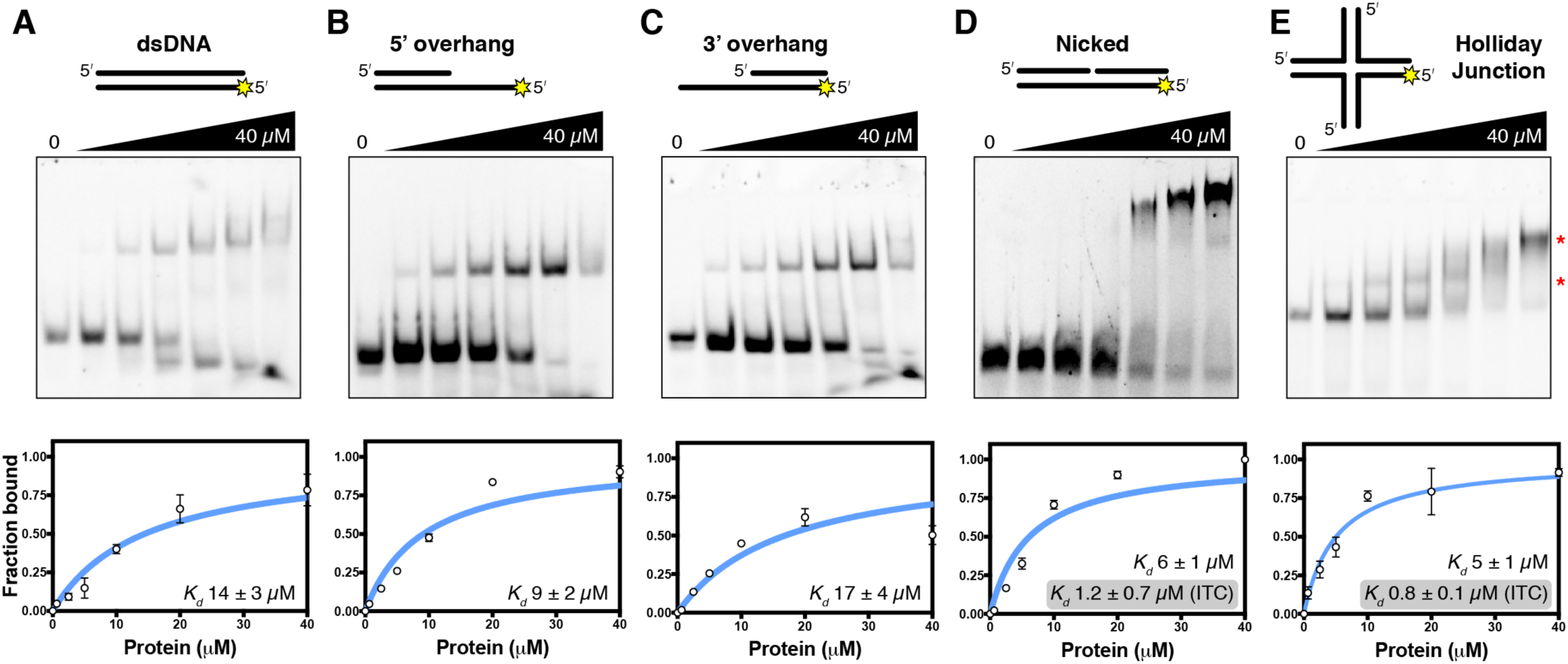
DNA binding by Zip2^499-704^:Spo16. (A-E) Representative gel-shift (*upper*) and binding curve from triplicate experiments (*lower*) for Zip2^499-704^:Spo16 binding dsDNA (A), 5’-overhang (B), 3’-overhang (C), nicked (D), and Holliday Junction (E) DNA substrates. Protein concentrations in each lane (left to right: 0, 0.625, 2.5, 5, 10, 20, and 40 μM) are the same for all gels. See **Figure S4A** for Zip2^499-704^:Spo16 binding to ssDNA. Red asterisks indicate location of multiple shifted bands for the HJ substrate. Grey boxes in panels (D) and (E) indicate *K*_*d*_values for nicked and Holliday Junction DNA as measured by isothermal titration calorimetry (**Figure S4B, C**).

We next used isothermal titration calorimetry (ITC) to verify binding of Zip2^499-704^:Spo16 to both the nicked and Holliday Junction substrates. Both substrates bound with ∽1 μM affinity in this assay, validating the results above and suggesting that our EMSA assays may systematically under-estimate the binding affinity of Zip2^499-704^:Spo16 for DNA (**Figure S4B,C**). Our ITC data also showed very slow equilibration kinetics and a large positive entropy change (*ΔS*) upon binding, suggesting that Zip2^499-704^:Spo16 may undergo a significant conformational change upon DNA binding.

Overall these data show that Zip2^499-704^:Spo16 robustly binds multiple DNA structures that likely exist in early meiotic recombination intermediates, including branched structures and ssDNA-dsDNA junctions with a 5’ overhang. The preferential binding of nicked and HJ DNA in particular over double-stranded or ssDNA-dsDNA junctions suggests that strongly bent or branched structures may be preferred substrates of the complex. Our data mirror recent findings by De Muyt et al., who found that a reconstituted Zip2(XPF domain):Spo16 complex binds branched DNA structures with higher affinity than dsDNA (De Muyt *et al*, 2018). Interestingly, De Muyt et al. identified a D-loop as the most-preferred substrate, but given the complexity of this substrate, which includes both ssDNA-dsDNA junctions and branched structures, it is unclear which element in this substrate is specifically recognized by Zip2:Spo16 (De Muyt *et al*, 2018). Similarly, Guiraldelli et al. recently showed that purified mammalian SHOC1 binds both D-loop and Holliday Junction structures more tightly than dsDNA, though these assays also included an N-terminal SHOC1 domain predicted to adopt a helicase fold, and did not include an ERCC1-like binding partner (Guiraldelli *et al*, 2018).

To identify likely DNA-binding surfaces on Zip2^499-704^:Spo16, we examined known DNA co-crystal structures of related enzymes. The *Aeropyrum pernix* XPF homodimer has been co-crystallized with double-stranded DNA (Newman *et al*, 2005), and the isolated HhH_2_domain of human XPF has been co-crystallized with single-stranded DNA (Das *et al*, 2012). In addition, human MUS81:EME1 has been co-crystallized with several more complex DNA structures including 5’ and 3’ flap structures (Gwon *et al*, 2014). The MUS81:EME1-DNA structures are particularly informative, as they show how these proteins’ central/nuclease and HhH_2_domains cooperate to bind a single DNA substrate and orient it for cleavage. To model the DNA-bound conformation of Zip2^499-704^:Spo16, we used a structure of MUS81:EME1 bound to a 5’-flap DNA substrate, which closely mimics the nicked DNA that Zip2^499-704^:Spo16 preferentially binds. The 5’-flap DNA is bent nearly 90° in the protein:DNA complex, in agreement with our speculation that bent or highly bendable structures (which would also include branched structures like Holliday Junctions) are preferred substrates of Zip2^499-704^:Spo16. In this structure, the central and HhH_2_ domains of MUS81:EME1 are separated, and sandwich the DNA substrate between them (Gwon *et al*, 2014). While Zip2^499-704^:Spo16 shows a consistent overall structure in all five views of the complex, closer inspection of the interface between the central and HhH domain dimers reveals that it is relatively small and mostly hydrophilic (not shown). Further, the inter-domain linkers on both Zip2 and Spo16 are long enough to allow significant conformational changes, and our ITC data suggested that the complex likely undergoes a conformational change upon binding DNA (**Figure S4B,C**). Thus, we separately overlaid the central and HhH domains of Zip2^499-704^:Spo16 onto the structure of MUS81:EME1. The resulting model shows a significant conformational shift of the HhH domains relative to their orientation in our crystal structures, but the two regions remain close enough that the inter-domain linkers in each protein can easily span the distance (**Figure 4A**). Based on this model, sequence alignments, and known DNA-binding surfaces of XPF-family proteins, we identified four surfaces on the Zip2^499-704^:Spo16 complex that are potentially involved in binding DNA. Patch 1, located on the Zip2 central domain, is close to the active site in related XPF-family proteins, and comprises three positively-charged residues: R554, R559, and K568 (**Figure 2E, 4B**). Patch 2 borders patch 1, comprising K600 of Zip2 and two Spo16 residues, R127 and N132 (**Figure 4B**). Both patches 1 and 2 are juxtaposed to DNA in our model. Patches 3 and 4 are located on adjacent surfaces of the Zip2 HhH_2_ domain, with patch 3 comprising residues K663, K683, and K686, and patch 4 comprising residues N657, Q692, and R695 (**Figure 4C**). Patch 3 is equivalent to a surface on human XPF previously shown to bind single-stranded DNA (Das *et al*, 2012), while patch 4 corresponds to the surface on MUS81 that interacts with double-stranded DNA (Gwon *et al*, 2014).

**Figure 4.**
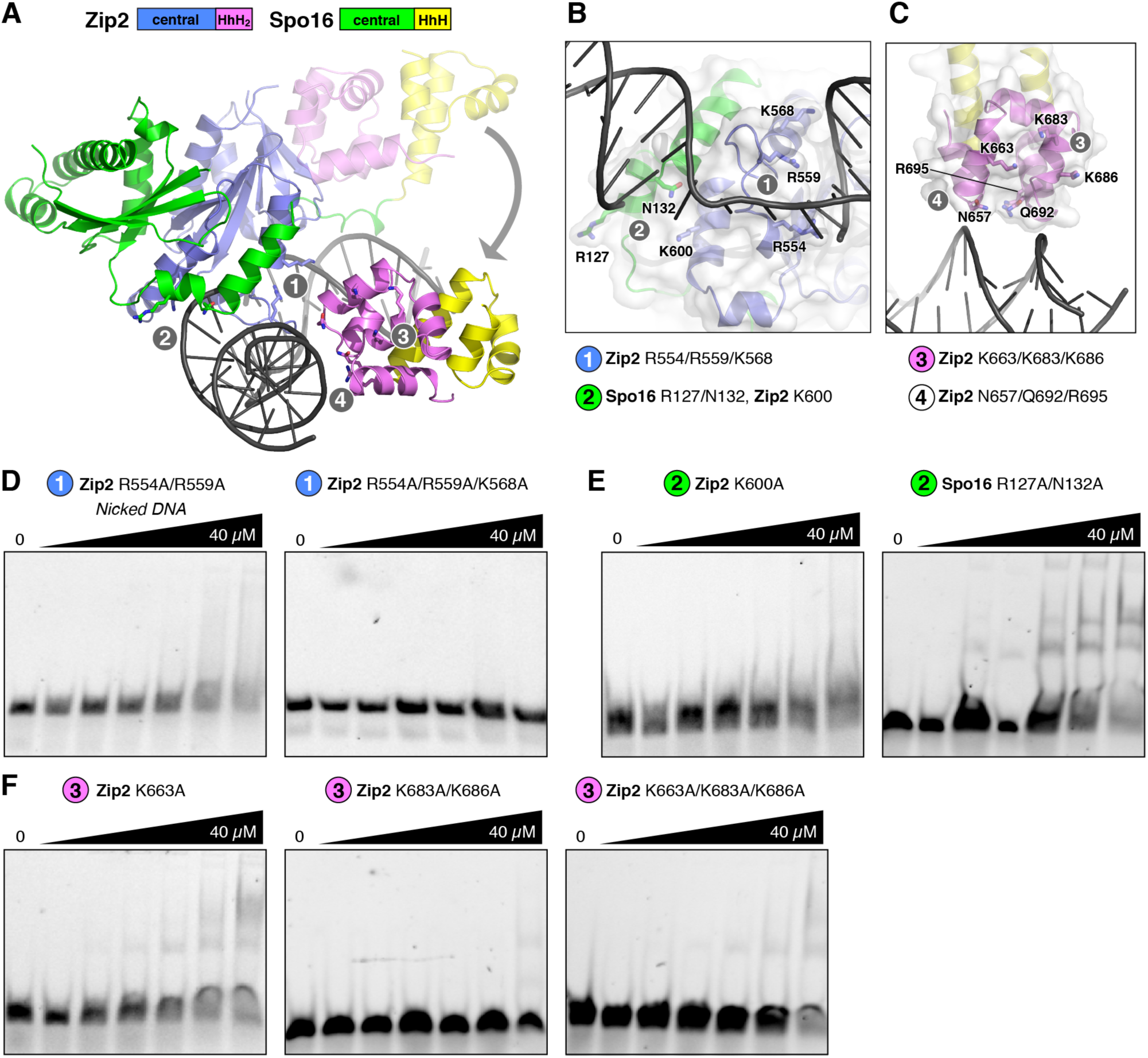
Identification of DNA-binding surfaces of Zip2^499-704^:Spo16. (A) Model of the Zip2^499-704^:Spo16 complex bound to a nicked/overhang DNA, based on a structure of human Mus81:Eme1 in complex with a 5’-flap DNA (PDB ID 4P0P) (Gwon *et al*, 2014). The central domains are oriented equivalently to **Figure 2A**. In the model, the HhH domains of Zip2 and Spo16 have undergone a significant conformational change from their original orientation (semi-transparent ribbons) upon DNA binding (gray arrow). Gray circles indicate the locations of putative DNA-binding patches 1-4. See **Figure S5A** for surface charge distribution. (B) Close-up view of patches 1 and 2 on the central domains of Zip2 and Spo16. See **Figure S5B** for surface charge distribution. (C) Close-up view of patches 3 and 4 on the Zip2 HhH_2_domain. See **Figure S5C** for surface charge distribution. (D) Representative gel-shift assays for patch 1 mutants binding nicked DNA. See **Figure S6A** for binding of 5’ overhang DNA. (E) Representative gel-shift assays for patch 2 mutants binding nicked DNA. See **Figure S6B** for binding of 5’ overhang DNA. (F) Representative gel-shift assays for patch 3 mutants binding nicked DNA. See **Figure S6C** for binding of 5’ overhang DNA. Patch 4 mutants were insoluble in vitro, precluding their analysis.

To test the roles of the four putative DNA-binding surfaces on Zip2^499-704^:Spo16, we mutated these surfaces and tested binding of both 5’ overhang and nicked DNA substrates by EMSA. We found that mutants in patches 1, 2, and 3 strongly affect binding to both DNA substrates, supporting our model of DNA binding by Zip2^499-704^:Spo16. Further, these results show that both the central and HhH domains, acting together, are required for robust DNA binding. We were unable to purify complexes containing mutants of patch 4, precluding their analysis.

## Discussion

The data we present here shows that the *S. cerevisiae* Zip2:Spo16 complex forms an XPF:ERCC1-like dimer with structure-specific DNA binding activity. We further show that the Zip2 N-terminal domain binds Zip4, scaffolding the assembly of a larger Zip2:Zip4:Spo16 complex. Based on prior work, the Zip2:Zip4:Spo16 complex is required for formation of type 1 interfering COs in yeast, and also is required for polymerization of the synaptonemal complex in coordination with crossover formation. Based on its structure and DNA binding activity, we propose that Zip2:Spo16 binds a specific DNA structure generated early in the recombination pathway, likely within the D-loop intermediate formed after initial strand invasion. This intermediate contains a ssDNA-dsDNA junction with a 5’ single-strand overhang, as well as a multi-strand junction equivalent to our nicked DNA substrate plus single-stranded flaps (**Figure 5**). Through this DNA binding activity, Zip2:Spo16 likely complements the activity of the Msh4:Msh5 complex, which is proposed to bind and stabilize a pre-Holliday Junction intermediate. Thus, Zip2:Spo16 and Msh4:Msh5 could together recognize and stabilize multiple structural elements of an early meiotic recombination intermediate, thereby stabilizing it against dissolution by topoisomerases/helicases to promote the crossover fate (**Figure 5**).

**Figure 5.**
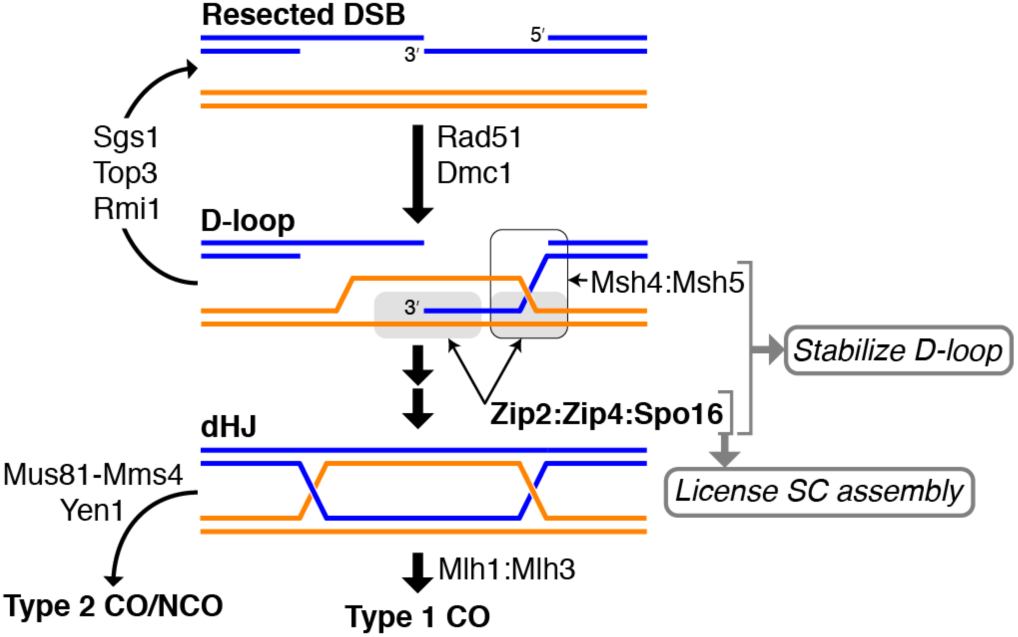
Model for the role of Zip2:Spo16:Zip4 in meiotic crossover formation. Model for the function of Zip2:Zip4:Spo16 in meiotic recombination. Initial strand invasion is mediated by Rad51 and Dmc1, and is continually counteracted by the dissolution activities of Sgs1, Top3, and Rmi1, resulting in SDSA. Recognition of specific DNA structures by Msh4:Msh5 (pro-HJ; black outline) and Zip2:Spo16 (ssDNA/dsDNA junction or pro-HJ; gray boxes) stabilize the strand – invasion/D-loop intermediate, recruit Type 1 CO-specific factors to promote the crossover outcome, and license assembly of the synaptonemal complex.

In addition to promoting and stabilizing early recombination intermediates, Zip2:Zip4:Spo16 is also important for assembly of the synaptonemal complex, likely linking events at the DNA level with “licensing” of synaptonemal complex assembly once recombination has progressed to a certain point. We propose that, similar to the role of FANCM:FAAP24 in the FA core complex, Zip2:Zip4:Spo16 can recruit and organize different proteins to promote the crossover fate and link crossover progression with chromosome axis remodeling and synaptonemal complex assembly. This recruitment is likely accomplished mostly by Zip4, which is predicted to fold into an array of 22 TPR repeats (Perry *et al*, 2005). Recently, De Muyt et al. identified a large number of potential interaction partners of the Zip2:Zip4:Spo16 complex using mass spectrometry, and demonstrated that Zip4 interacts directly with several of these, including the axis protein Red1 and the ZMM proteins Zip3 and Msh5 (De Muyt *et al*, 2018). These data strongly support a model in which Zip2 and Spo16 recognize and stabilize a specific early recombination intermediate, while Zip4 is largely responsible for specific protein-protein interactions necessary for linking DNA recognition to downstream events including later stages of recombination and chromosome morphology changes.

In addition to protein interactions mediated by Zip4, close inspection of our Zip2^499-704^:Spo16 complex reveals a conserved, concave hydrophobic surface involving the C-terminal W704 residue of Zip2 and two *α*-helices from the Spo16 HhH domain (**Figure S7**). This interface binds a short hydrophobic *α*-helix on the Zip2 central domain in three of the five independent views of the complex (two of the four copies in Form 1, plus the single copy from Form 2), burying 410 Å^2^ of mostly-hydrophobic surface area on each partner. Based on the high conservation of the involved residues on the Spo16 HhH domain, their mostly hydrophobic nature, and our observation of this interface in both crystal forms of Zip2^499-704^:Spo16, we propose that this surface may be involved in specific protein-protein interactions. While our initial yeast two-hybrid assays revealed no interactions between Zip2:Spo16 and other known ZMM/chromosome axis/synaptonemal complex proteins (not shown), the role of this surface in protein-protein interactions will be an interesting avenue for future work, especially given the likelihood that Zip2:Zip4:Spo16 acts a scaffolding complex for DNA repair and synaptonemal complex assembly.

In *A. thaliana*, SHOC1 and PTD interact with one another and are required for the formation of type 1 COs, and sequence analyses have suggested that these proteins also form an XPF:ERCC1-like dimer (Macaisne *et al*, 2011; 2008). More recent work has shown that human SHOC1 localizes to meiotic recombination sites and is required for MLH1 localization and formation of chiasmata (Guiraldelli *et al*, 2018). When considered alongside the key roles of Zip2:Zip4:Spo16 in *S. cerevisiae* meiosis, these data strongly support the idea that an XPF:ERCC1-like DNA binding complex is an important and highly conserved feature of meiotic crossover control in eukaryotes. Further work will be required to identify Zip4-like scaffolding subunits in organisms outside fungi, and to determine how this conserved complex cooperates with Msh4:Msh5 to promote the crossover fate, and coordinate recombination with chromosome morphology changes such as chromosome axis remodeling and synaptonemal complex assembly.

## Materials and Methods

For key resources, see **Table S2**.

### Yeast two-hybrid assays

For yeast two-hybrid assays, individual proteins were cloned into pBridge and pGADT7 AD vectors (Clontech) with multiple-cloning sites modified for ligation-independent cloning (http://qb3.berkeley.edu/macrolab/lic-cloning-protocol/). pBridge vectors were transformed into *S. cerevisiae* strain AH109 and selected on SC media lacking tryptophan (-TRP). pGADT7 AD vectors were transformed into *S. cerevisiae* strain Y187 and selected on SC media lacking leucine (-LEU). Haploid strains were mated and diploids selected on SC -TRP/-LEU. Diploid cells were diluted in water and replated onto SC-TRP/-LEU (control),-TRP/-LEU/-HIS (histidine) (low stringency), and-TRP/-LEU/-HIS/-ADE (adenine) (high stringency), grown for 2-3 days, then examined for growth.

For yeast three-hybrid assays, pBridge vectors containing either Spo16 or Zip2 in MCS I were further modified by NotI cleavage at the MCS II site followed by isothermal assembly-mediated insertion of the second gene (Gibson *et al*, 2009), resulting in a single vector encoding a Zip2:Spo16 complex containing the Gal4-BD tag fused to the N-terminus of either protein. These vectors were transformed into AH109 and mated with pGADT7 AD vectors encoding Zip4.

### Protein expression and purification

Ligation-independent cloning was used to clone full length Spo16 and Zip2^499-704^. To express TEV protease-cleavable, His_6_-tagged Spo16, full-length Spo16 was cloned into Addgene vector 48324 (contains Spectinomycin resistance and CloDF13 ori) using ligation-independent cloning. To express untagged Zip2 499-704, DNA encoding Zip2 499-704 was cloned into Addgene vector 29665 (contains Ampicillin resistance and ColE1 origin of replication) using ligation-independent cloning. The Zip2 surface entropy reduction (SER) mutant (KEK 641-643 → AAA) was identified by the UCLA SERp server (http://services.mbi.ucla.edu/SER/) (Goldschmidt *et al*, 2007) and was generated by mutagenic PCR.

For protein expression, plasmids encoding Zip2^499-704^ (unmutated or SER mutant) and full-length Spo16 were co-transformed into *E. coli* Rosetta2 (DE3) pLysS cells, and grown in 2XYT media supplemented with both ampicillin and spectinomycin. Cells were grown at 37°C to and OD_600_of 0.6, shifted to 20°C and protein expression induced with 0.25 mM IPTG, and grown 16 hours. For selenomethionine derivatization, cells were grown in M9 minimal media at 37°C to an OD_600_of 0.8, after which the following amino acids were added: Leu, Ile and Val (50 mg/L), Phe, Lys, Thr (100 mg/L) and Selenomethionine (60 mg/L). Cells were shifted to 20°C and protein expression was induced with 0.25 mM IPTG after 20 minutes incubation with amino acids.

For protein purification, cells were harvested by centrifugation, suspended in resuspension buffer (20 mM Tris-HCl pH 8.0, 300 mM NaCl, 10 mM imidazole, 1 mM dithiothreitol (DTT) and 10% glycerol) and lysed by sonication. Lysate was clarified by centrifugation (16,000 rpm 30 min), then supernatant was loaded onto a Ni^2+^ affinity column (HisTrap HP, GE Life Sciences) pre-equilibrated with resuspension buffer. The column was washed with buffer containing 20 mM imidazole and 100 mM NaCl, and eluted with a buffer containing 250 mM imidazole and 100 mM NaCl. The elution was loaded onto an anion-exchange column (Hitrap Q HP, GE Life Sciences) and eluted using a 100-600 mM NaCl gradient. Fractions containing the protein were pooled and concentrated to 2 mL by ultrafiltration (Amicon Ultra-15, EMD Millipore), then passed over a size exclusion column (HiLoad Superdex 200 PG, GE Life Sciences) in a buffer containing 20 mM Tris-HCl pH 8.0, 200 mM NaCl, and 1 mM DTT. The N-terminal His_6_-tag on Zip2, while cleavable by TEV protease, was not removed for any experiments shown. Purified proteins were concentrated by ultrafiltration and stored at 4°C for crystallization, or aliquoted and frozen at-80°C for biochemical assays.

Surface lysine residues on the Zip2^499-704^:His_6_-Spo16 complex were dimethylated by mixing protein at 1 mg/mL with 20 mM freshly-prepared Dimethylamine Borane Complex and 40 mM formaldehyde, incubation at 4°C for one hour, then quenching by addition of 100 mM glycine. Alkylated proteins were concentrated and purified by size-exclusion chromatography as above. Dimethyl-lysine residues can be observed in several positions in electron density maps of Form 1 crystals (**Figure S3A, B**).

For size exclusion chromatography coupled to multi-angle light scattering (SEC-MALS), 100 μL of Zip2^499-704^:His_6_-Spo16 at 2.0 mg/mL was injected onto a Superdex 200 Increase 10/300 GL column (GE Life Sciences) in a buffer containing 20 mM HEPES pH 7.5, 300 mM NaCl, 5% glycerol, and 1 mM DTT. Light scattering and refractive index profiles were collected by miniDAWN TREOS and Optilab T-rEX detectors (Wyatt Technology), respectively, and molecular weight was calculated using ASTRA v. 6 software (Wyatt Technology).

### Protein crystallization

Form 1 crystals were obtained in hanging drops with surface-lysine methylated Zip2^499-704^:His_6_-Spo16 at 10 mg/mL in a buffer containing 20 mM Tris-HCl pH 7.5 and 200 mM NaCl. Protein was mixed 1:1 with well solution containing 1.4 M Na-K phosphate pH 6.6. Crystals were transferred to a cryoprotectant solution containing 2.4 M Na malonate pH 7.0, then flash-frozen in liquid nitrogen. Form 2 crystals were obtained in hanging drops with Zip2^499-704^SER:His_6_-Spo16 at 10 mg/mL in a buffer containing 20 mM Tris-HCl pH 7.5 and 200 mM NaCl. Protein was mixed 1:1 with well solution containing 100 mM Bis Tris pH 5.5, 200 mM Ammonium sulfate, 15% PEG 3350. Crystals were cryo-protected by addition of 15-20% PEG 400, then flash-frozen in liquid nitrogen.

### X-Ray data collection and structure determination

For Form 1, diffraction data was collected at the Advanced Photon Source, NE-CAT beamline 24ID-E (support statement below). Data was automatically indexed and reduced by the RAPD data-processing pipeline (https://github.com/RAPD/RAPD), which uses XDS (Kabsch, 2010) for indexing and integration, and the CCP4 programs AIMLESS and TRUNCATE (Winn *et al*, 2011) for scaling and structure-factor calculation. Extensive efforts to determine the structure by anomalous methods using selenomethionine-derivatized protein failed due to translational pseudo-symmetry arising from the positions and orientations of the four copies of Zip2^499-704^:Spo16 in these crystals.

For Form 2, diffraction data for both native and selenomethionine-derivatized crystals were collected at the Advanced Photon Source, NE-CAT beamline 24ID-C, and data was automatically indexed and reduced by RAPD. The structure was determined by single-wavelength anomalous diffraction (SAD) methods using a 2.38 Å-resolution dataset collected from selenomethionine-derivatized proteins. Selenium sites were located using hkl2map/SHELX (Pape *et al*, 2004; Sheldrick, 2010), and provided to the Phenix Autosol pipeline (Terwilliger *et al*, 2009; Adams *et al*, 2010) for phase calculation using PHASER (McCoy *et al*, 2007) and density modification using RESOLVE (Terwilliger, 2000). A partial model built by RESOLVE (Terwilliger, 2003) was manually rebuilt in COOT (Emsley *et al*, 2010) and refined against a 2.13 Å-resolution native dataset using phenix.refine (Afonine *et al*, 2012).

To determine the Form 1 structure, molecular replacement was performed using PHASER to place four copies of the Form 1 dimer structure. The model was manually rebuilt in COOT and refined in phenix.refine to 2.29 Å resolution. Refined electron-density maps revealed additional electron density extending the side-chain of Zip2 Cys521 in all four copies of the complex. Formaldehyde, which was used for surface lysine alkylation of the complex, can react with the cysteine side-chain to produce hydroxymethylcysteine (Bateman *et al*, 2007; Metz *et al*, 2004), which closely matches the observed electron density for this residue (**Figure S3C,D**). We also observed clear electron density for several residues in the N-terminal His_6_-tag fused to Spo16, which included a TEV protease cleavage site (MKSSHHHHHHENLYFQ^SNA-[Spo16^2-198^]), packing against a symmetry-related copy of Spo16, explaining why these crystals required an intact tag for growth (**Figure S3E**). Data collection and refinement statistics for both structures can be found in **Table S1**. All structure figures were created with PyMOL version 2, and surface charge calculations were performed with the APBS (Jurrus *et al*, 2018) plugin in PyMOL. Original diffraction data have been deposited with the SBGrid Data Bank (https://data.sbgrid.org) under accession numbers 538 (Zip2^499-704^:Spo16 Form 1 Native), 539 (Zip2^499-704^SER:Spo16 Form 2 Native), and 540 (Zip2^499-704^SER:Spo16 Form 2, selenomethionine-derivatized SAD dataset). Reduced data and refined structures have been deposited with the RCSB Protein Data Bank (https://www.rcsb.org) under accession numbers 6BZF (Zip2^499-704^:Spo16 Form 1) and 6BZG (Zip2^499-704^SER:Spo16 Form 2).

### APS NE-CAT Support Statement

This work is based upon research conducted at the Northeastern Collaborative Access Team beamlines, which are funded by the National Institute of General Medical Sciences from the National Institutes of Health (P41 GM103403). The Pilatus 6M detector on 24-ID-C beam line is funded by a NIH-ORIP HEI grant (S10 RR029205). This research used resources of the Advanced Photon Source, a U.S. Department of Energy (DOE) Office of Science User Facility operated for the DOE Office of Science by Argonne National Laboratory under Contract No. DE-AC02-06CH11357.

### DNA binding assays

To generate different DNA substrates for electrophoretic mobility shift assays, a 40-base oligonucleotide 5’-labeled with 6-carboxyfluorescein (5’-6-FAM_40bp) was annealed at 10 μM concentration in annealing buffer (1X Tris-HCl pH 8.0, 50 mM NaCl, 1 mM MgCl_2_, 1 mM EDTA) with specific unlabeled oligos (sequences in **Table S2**) as follows: ssDNA (5’-6-FAM_40bp alone); dsDNA (5’-6-FAM_40bp + 40bp_for_ds); 5’-overhang (5’-6-FAM_40bp + 20bp_for_free5’); 3’-overhang (5’-6-FAM_40bp + 20bp_for_free3’); HJ (5’-6-FAM_40bp + HJ_strand2 + HJ_strand3 + HJ_strand4). Annealing was performed in a PCR machine using a temperature gradient from 95°C to 4°C, at a speed of 0.1°C per second. The EMSA reaction with 40-bp dsDNA or free 5’ overhang were prepared (buffer contained 20 mM Tris-HCl pH 8.0, 1 mM DTT, 5 mM MgCl2, 5% glycerol) by keeping the DNA concentration constant and varying the protein concentration. After 10 min incubation and addition of 5% (w/v) sucrose, free DNA and DNA-protein complexes were resolved by electrophoresis on 6% TBE-acrylamide gels pre-equilibrated (pre-run for 30 min at 150 V) in 0.2X TBE running buffer. Gels were run for 40 min at 100V at 4°C. Gels were imaged using a Bio-Rad ChemiDoc system using filters to image Cy2 dye. Gel bands were quantified using ImageJ (https://imagej.net), and binding curves were calculated using GraphPad Prism (https://www.graphpad.com) using a single-site binding model:

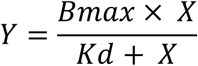

Where *Y* is fraction of DNA bound, *X* is protein concentration, *Bmax* is maximum possible DNA bound (constrained to 1), and *Kd* is the dissociation constant.

Isothermal titration calorimetry was performed on a Microcal ITC 200 (Malvern Panalytical) in a buffer containing 20 mM Tris (pH 8.5), 300 mM NaCl, and 1 mM TCEP. Zip2^499-704^:Spo16 at 230 μM was injected into an analysis cell containing 20 or 25 μM DNA.

## Acknowledgements

The authors thank the staff of NE-CAT Sector 24 beamlines at the Advanced Photon Source for assistance with data collection, processing, and phasing; A. Bobkov (Sanford Burnham Prebys Medical Discovery Institute, Protein Analysis Core Facility) for assistance with isothermal titration calorimetry; A. De Muyt and V. Borde for sharing information prior to publication; and members of the Corbett lab for helpful discussions. This work was supported by NIH/NIGMS grant number R01GM104141.

## Author Contributions

K.A. and K.D.C. designed the study. K.A. purified proteins, determined x-ray crystal structures, and performed DNA binding assays. K.A. and K.D.C. analyzed and interpreted data, and wrote the manuscript.

## Supplemental Figures

**Figure S1.**
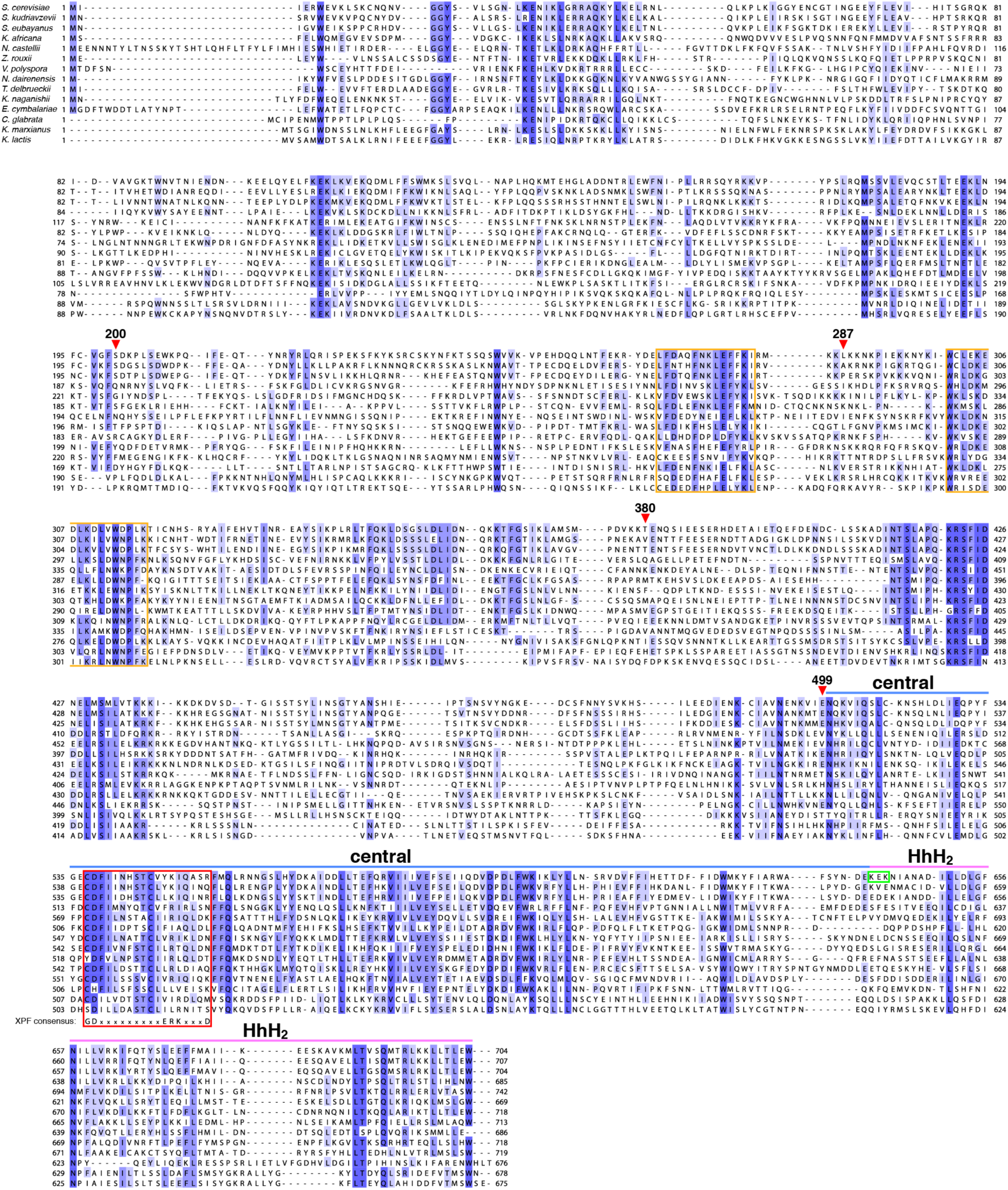
Sequence alignment of fungal Zip2 proteins. Red arrowheads indicate N-terminal truncations used for yeast two-and three-hybrid analysis. Boxed in orange are the conserved motifs spanning residues 269-282 and 301-317, potentially involved in Zip4 binding. Boxed in green is the motif (KEK, residues 641-643) mutated to alanine in the Zip2^499-704^SER mutant. Blue and violet lines indicate the extent of the central and HhH_2_domains.

**Figure S2.**
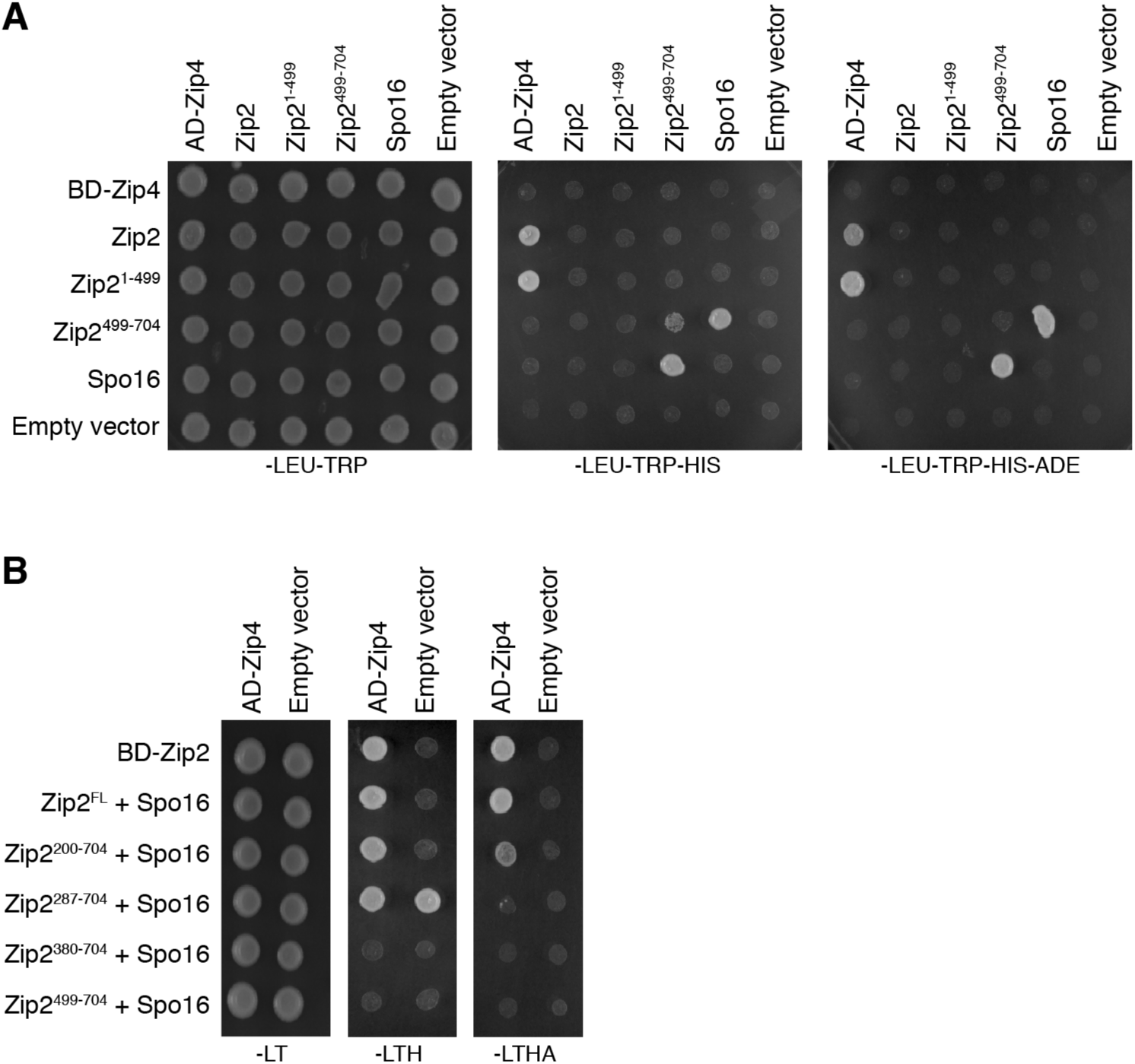
Yeast two-hybrid and three-hybrid analysis of Zip2, Zip4, and Spo16. (A) Yeast two-hybrid analysis of interactions between Zip2, Zip4, and Spo16. (B) Yeast three-hybrid analysis, with one vector (BD) encoding full-length or truncated Zip2 plus full-length Spo16, and the second vector (AD) encoding full-length Zip4.

**Figure S3.**
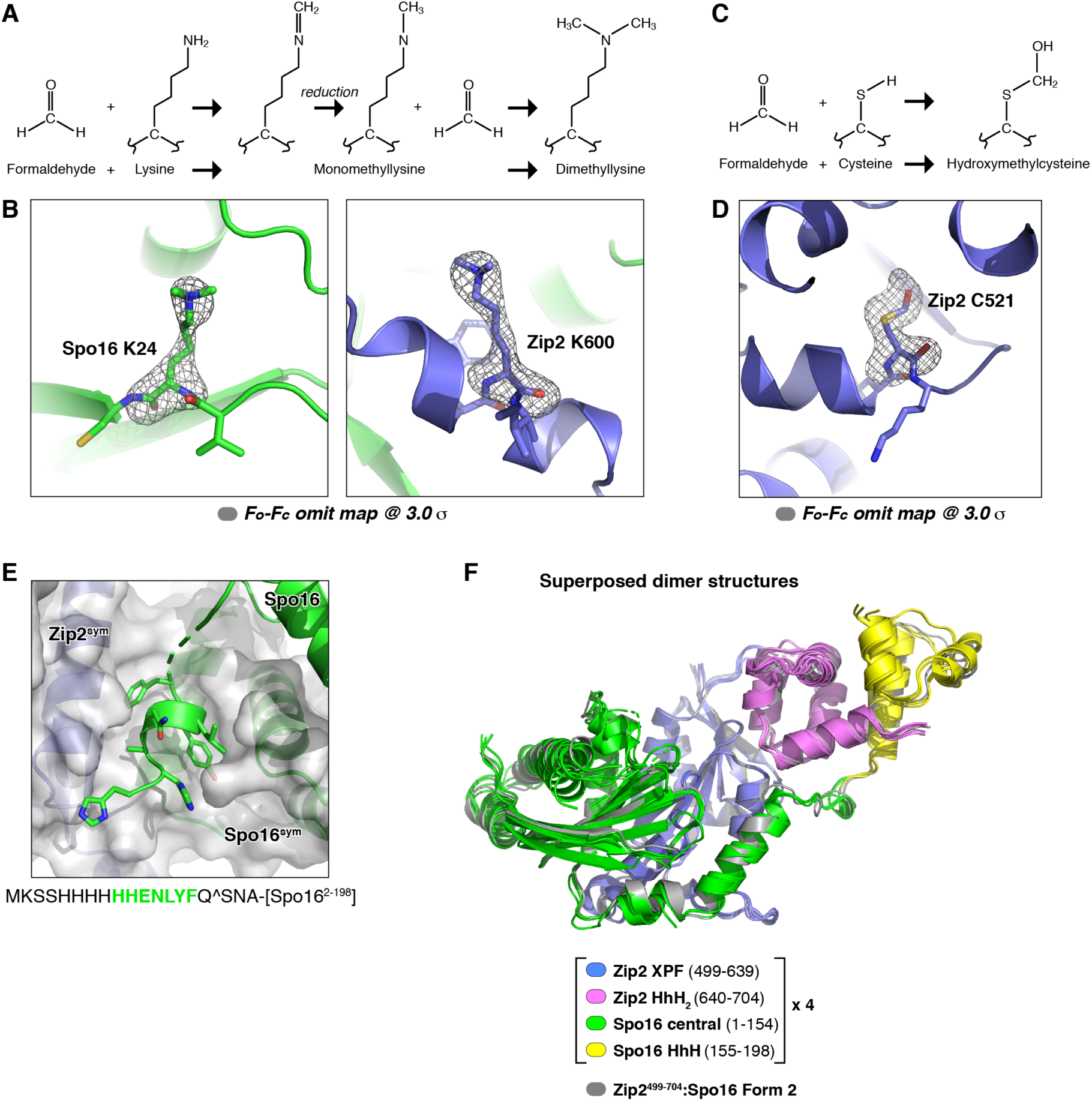
Reductive alkylation and crystal packing in Zip2^499-704^:Spo16 crystals. (A) Schematic for reductive methylation of surface lysine residues by formaldehyde. (B) Representative electron density (*F*_*o*_*-F*_*c*_map contoured at 3.0 *σ*, calculated from a model missing the modified residue) for two dimethyl-lysine residues. (C) Schematic for the formation of hydroxymethylcysteine by reaction of cysteine with formaldehyde. (D) Representative *F*_*o*_*-F*_*c*_ omit electron density (3.0 *σ*) for Zip2 cysteine 521. Chain A shown; the modified cysteine residue is clearly identifiable in all four Zip2 monomers of Form 1 crystals. (E) View of crystal packing interactions mediated by the N-terminal His_6_-tag on Spo16, which also contains a TEV protease cleavage site. Residues from the tag that are visible in electron density are highlighted in green. (F) Overlay of the five crystallographically-independent views of Zip2^499-704^:Spo16. The four copies of the complex in crystal form 1 are shown colored as in Figure 2, and the single copy of Zip2^499-704^ SER:Spo16 in crystal form 2 is shown in gray. Overall C*α* r.m.s.d. for all five copies ranges from 0.3 to 1.4 Å.

**Figure S4.**
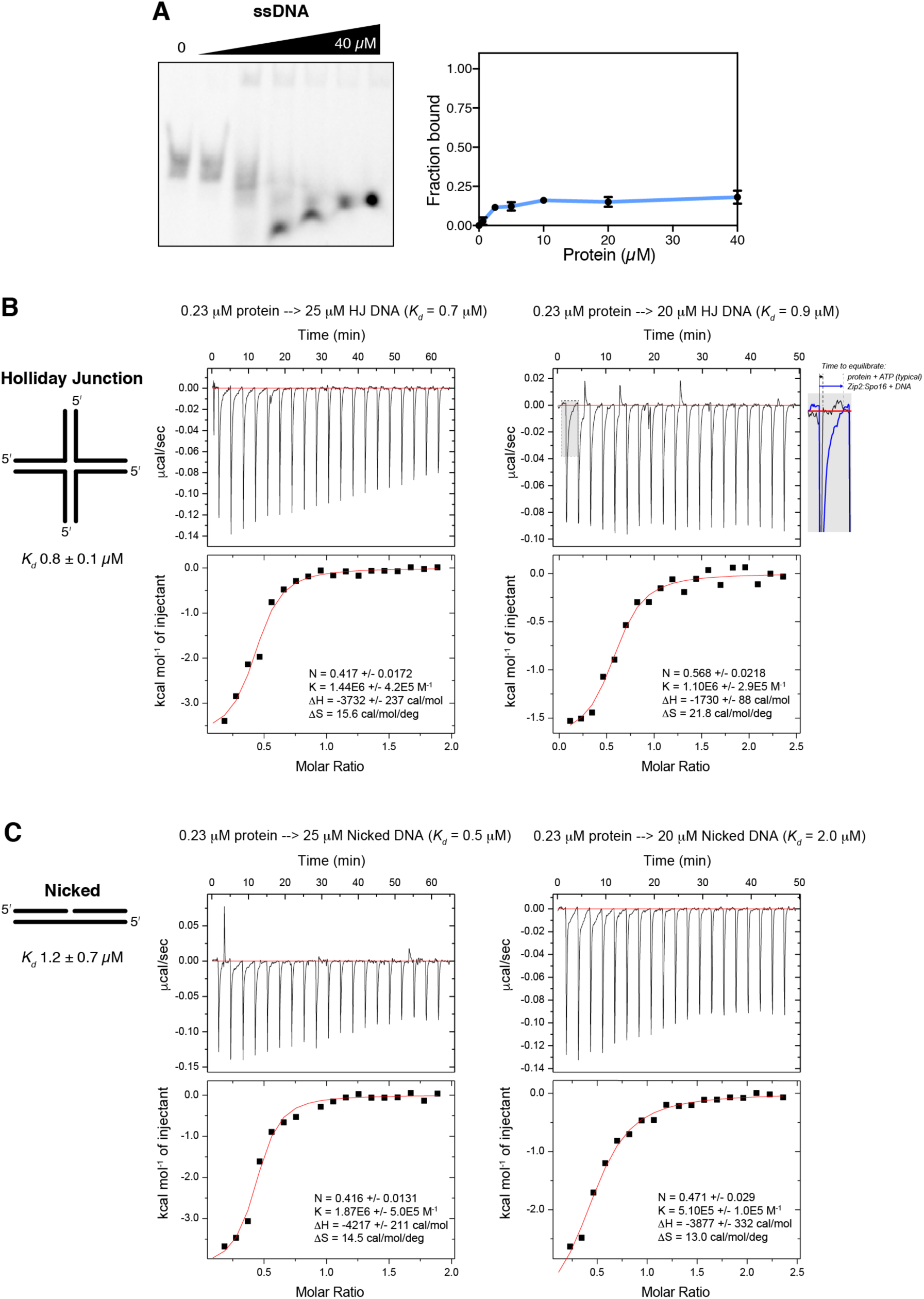
DNA binding by the Zip2^499-704^:Spo16 complex. (A) Representative gel-shift and binding curve for Zip2^499-704^:Spo16 binding single-stranded DNA. DNA-binding affinity was not calculated. (B) Isothermal titration calorimetry (ITC) for Zip2^499-704^:Spo16 binding to Holliday Junction DNA (duplicate experiments). Gray box and zoomed-in panel at far right show the slow equilibration for DNA binding (blue line) compared to a typical protein + ATP binding equilibration (gray line). The slow equilibration and large positive entropy change (*ΔS*) upon binding suggest that the protein complex undergoes a conformational change upon binding DNA. (C) Isothermal titration calorimetry (ITC) for Zip2^499-704^:Spo16 binding to nicked DNA (duplicate experiments).

**Figure S5.**
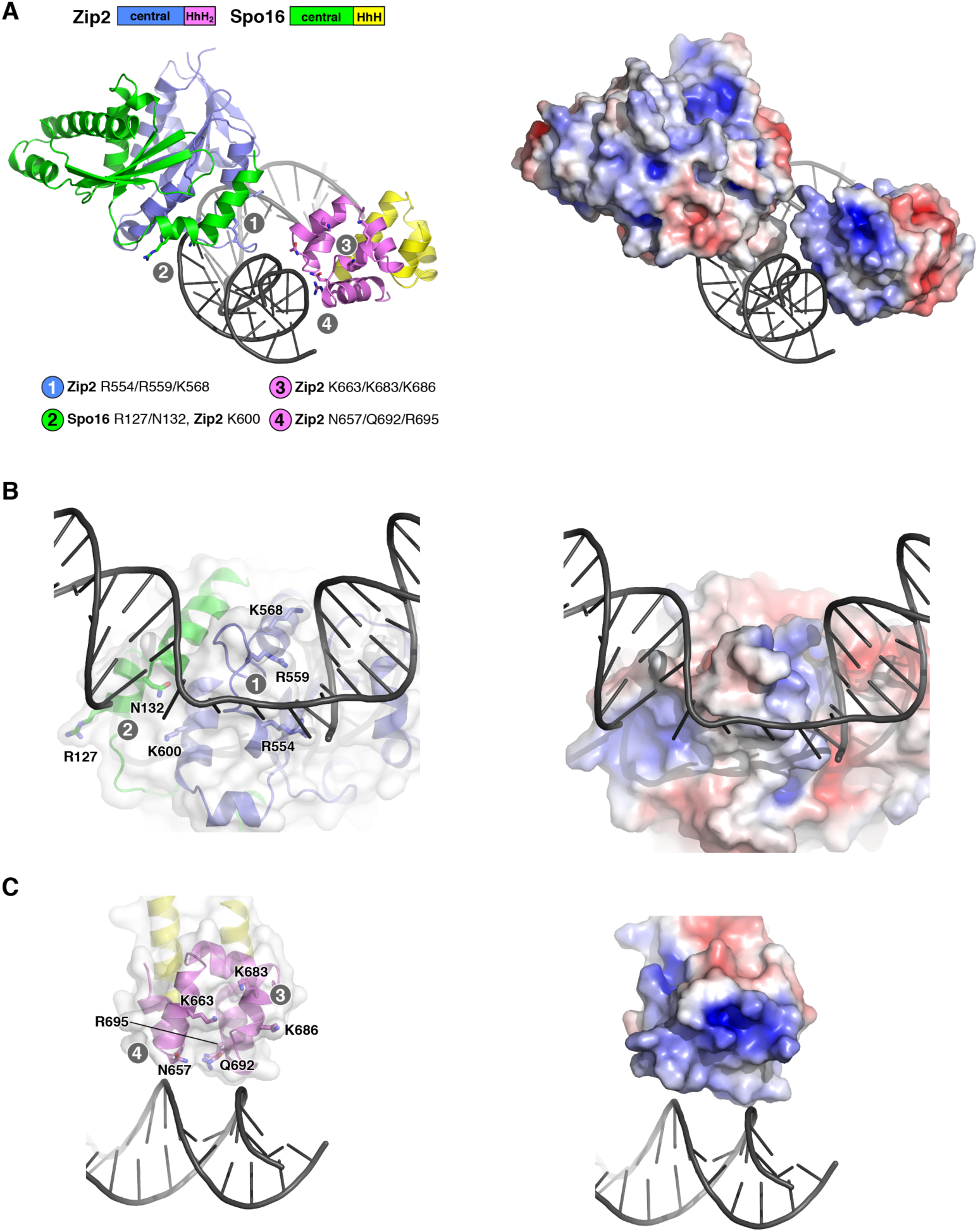
Identification of putative DNA-binding surfaces of Zip2^499-704^:Spo16. *(A) Left:* View equivalent to **Figure 4A**, showing a model of Zip2^499-704^:Spo16 bound to dsDNA, based on the structure of human MUS81:EME1 bound to a 5’-flap DNA (the ssDNA flap itself is disordered in the original structure) (PDB ID 4P0P; (Gwon *et al*, 2014)). The central and HhH regions were separately modeled, resulting in a distance of 40 Å between Zip2 residues 633 and 646 (this stretch can potentially span ∽50 Å, given 3.8 Å C*α*-C*α* distance for a fully-extended chain), and ∽31 Å between Spo16 residues 145 and 157 (which can span 46 Å). Putative DNA-binding patches 1-4 are denoted in gray. *Right:* Protein surface charge distribution of the modeled complex. *(B) Left:* Close-up view of patches 1 and 2 on the central domains of Zip2 and Spo16. *Right:* Protein surface charge distribution. *(C) Left:* Close-up view of patches 3 and 4 on the Zip2 HhH_2_ domain. *Right:* Protein surface charge distribution.

**Figure S6.**
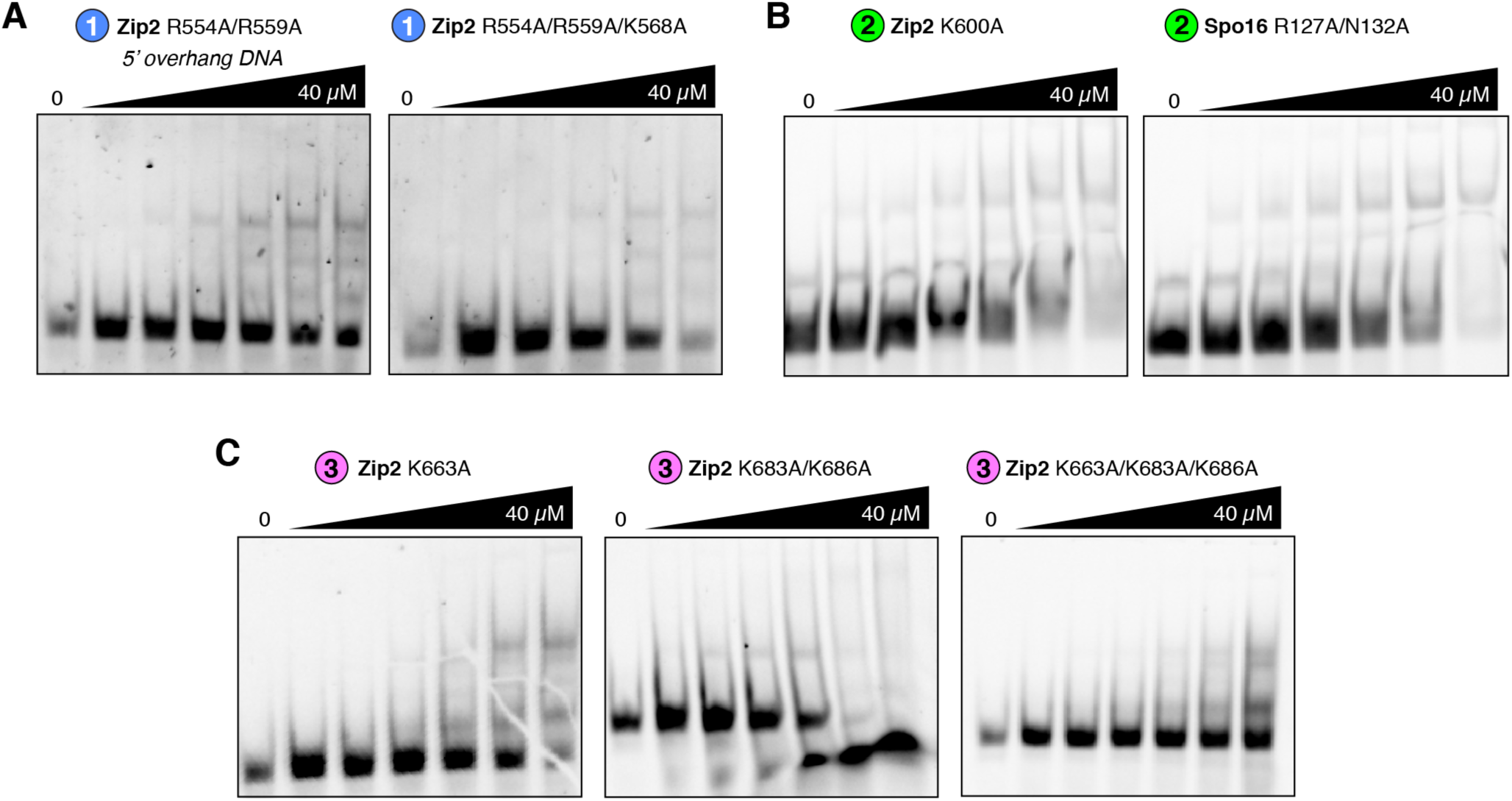
Binding of mutant Zip2:Spo16 complexes to 5’ overhang DNA. (A) Representative gel-shift assays for patch 1 mutants binding 5’ overhang DNA. (B) Representative gel-shift assays for patch 2 mutants binding 5’ overhang DNA. (C) Representative gel-shift assays for patch 3 mutants binding 5’ overhang DNA.

**Figure S7.**
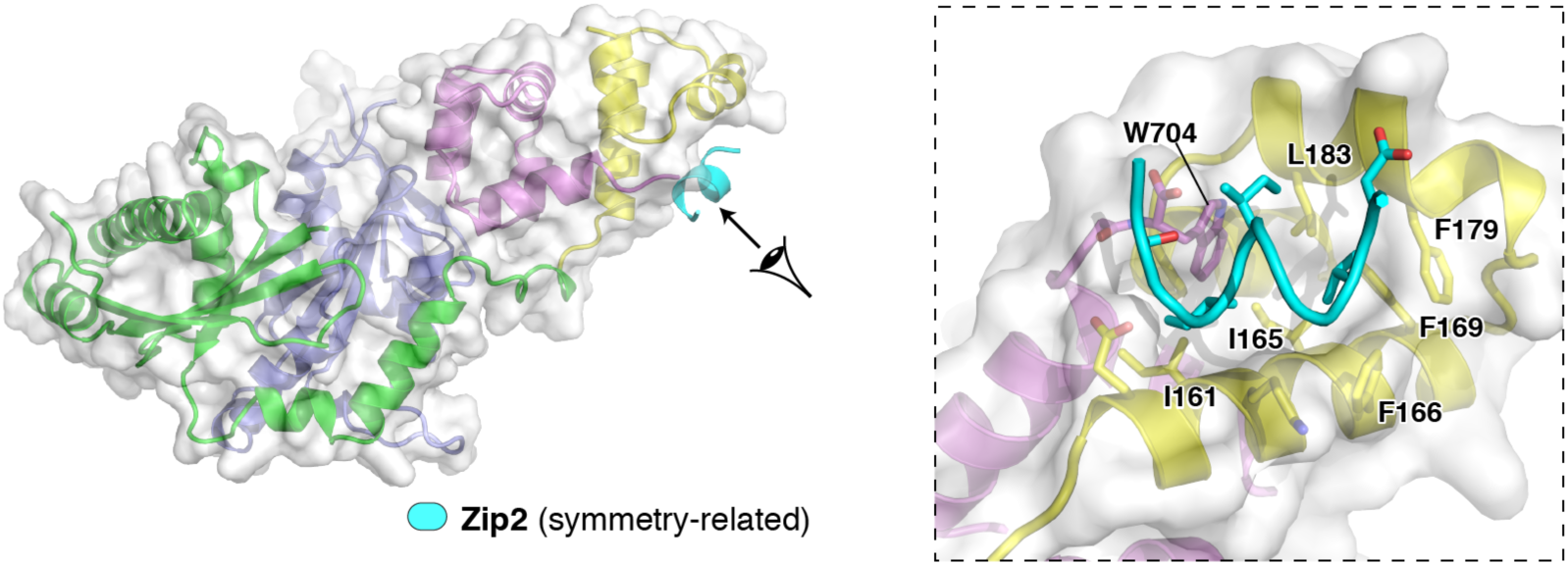
Identification of a putative protein-protein interaction surface on Spo16. Overall (left) and close-up (right) views of the Zip2^499-704^complex, with a crystallographic symmetry – related *α*-helix from Zip2 shown in cyan. This *α*-helix packs against a conserved hydrophobic cavity comprising the C-terminal W704 residue of Zip2 and several residues in the Spo16 HhH domain. This interface is observed in Form 2 crystals and two of the four dimers in Form 1 crystals.

**Table S1.**
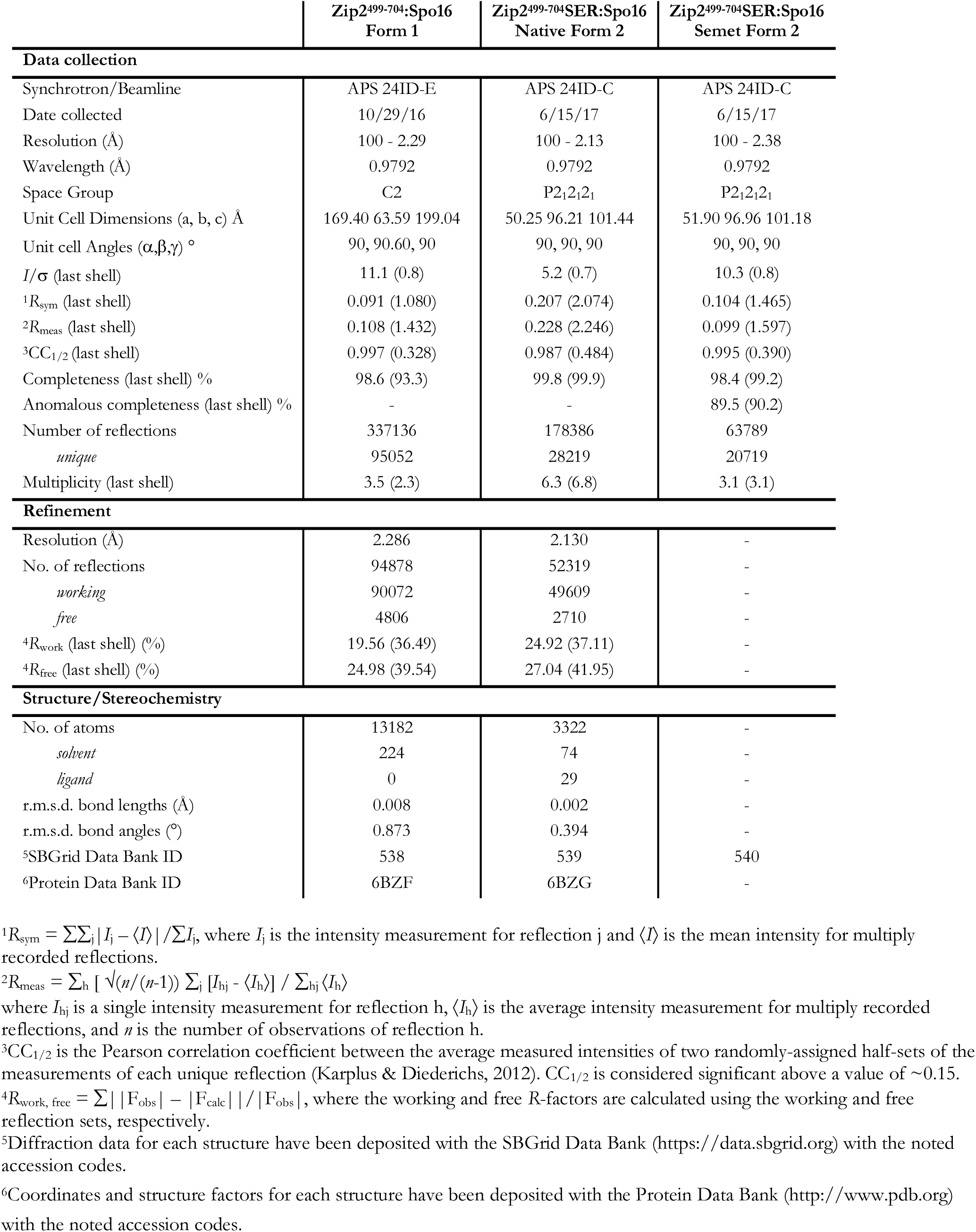
Crystallographic data and refinement

**Table S2.**
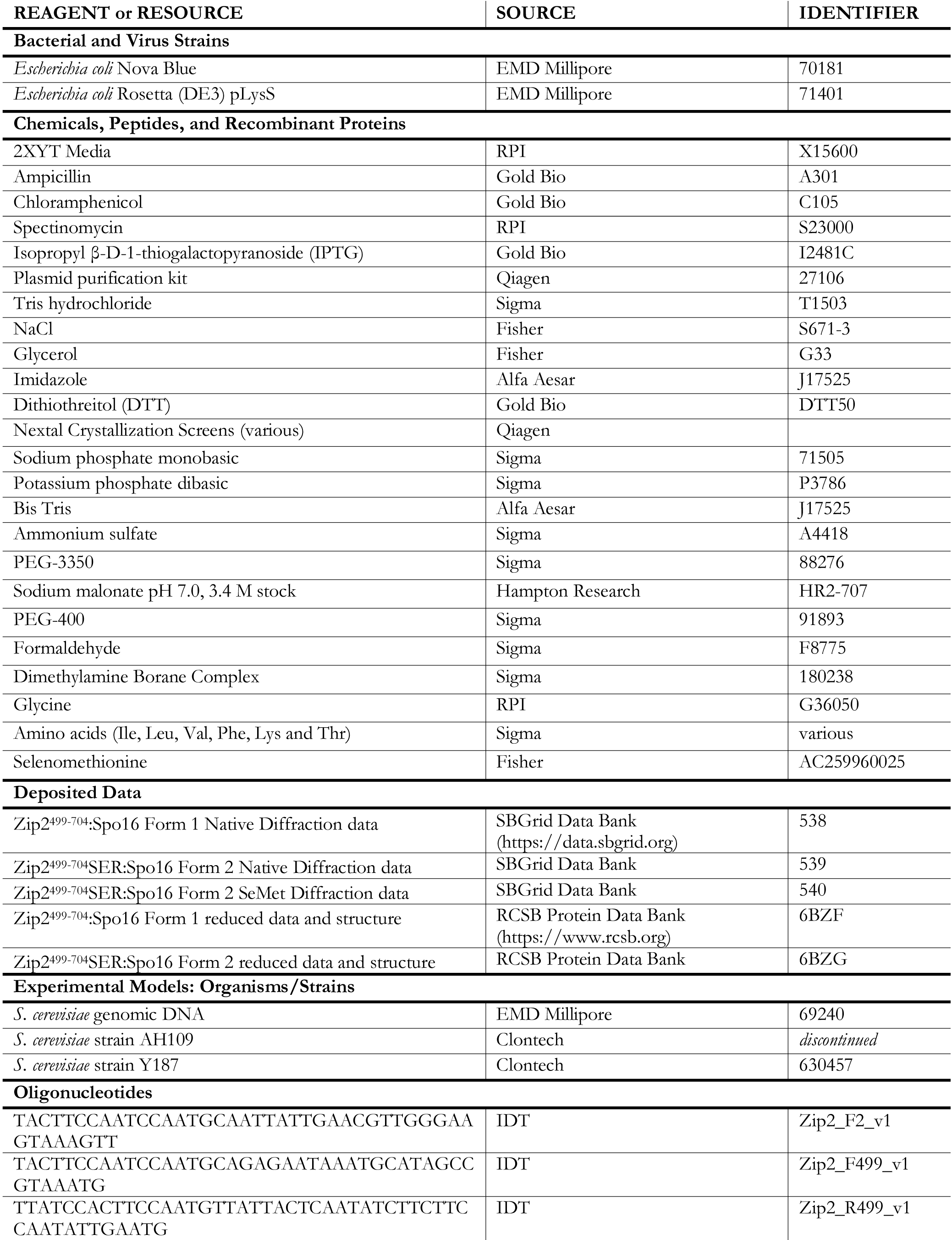
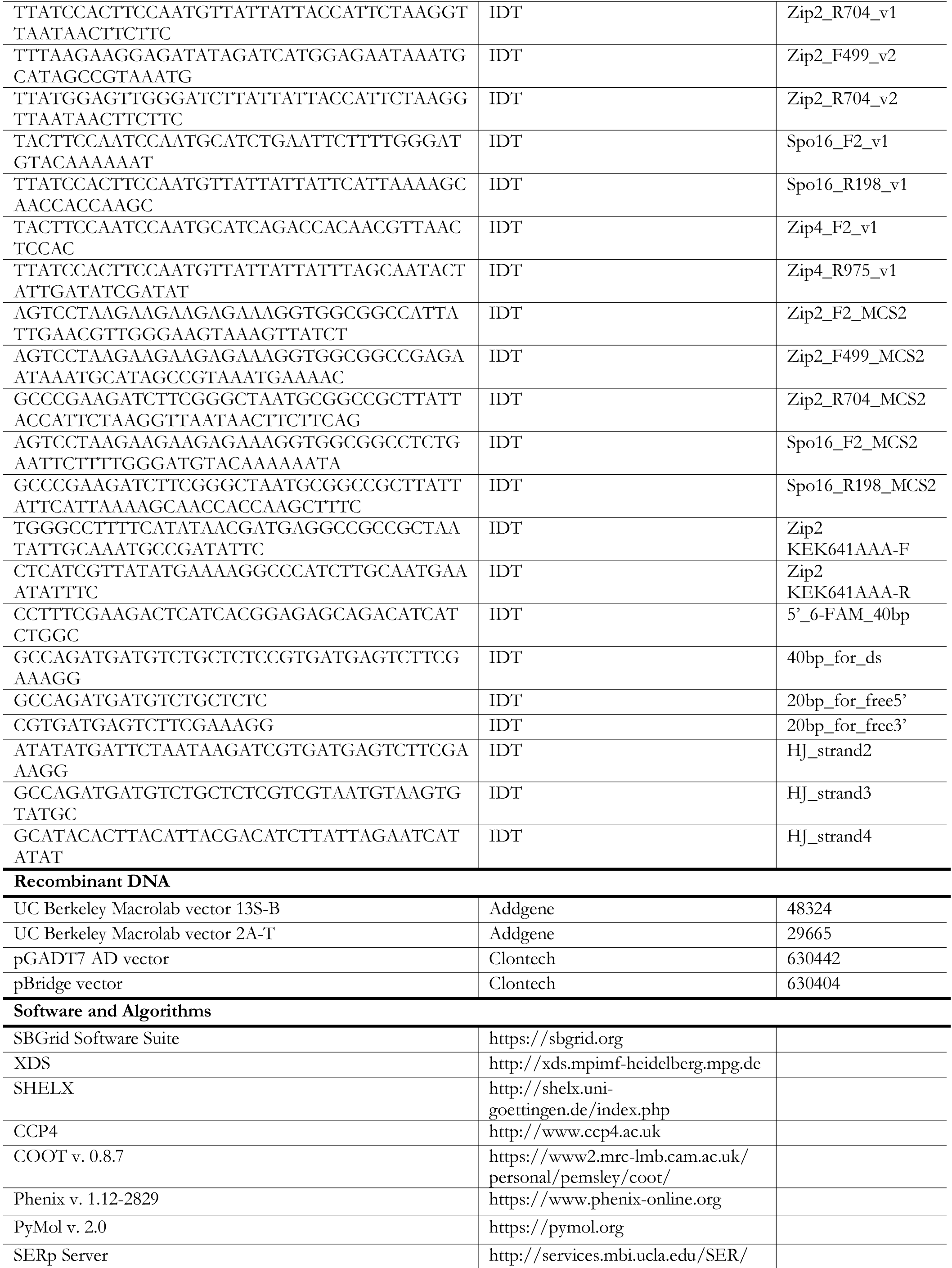
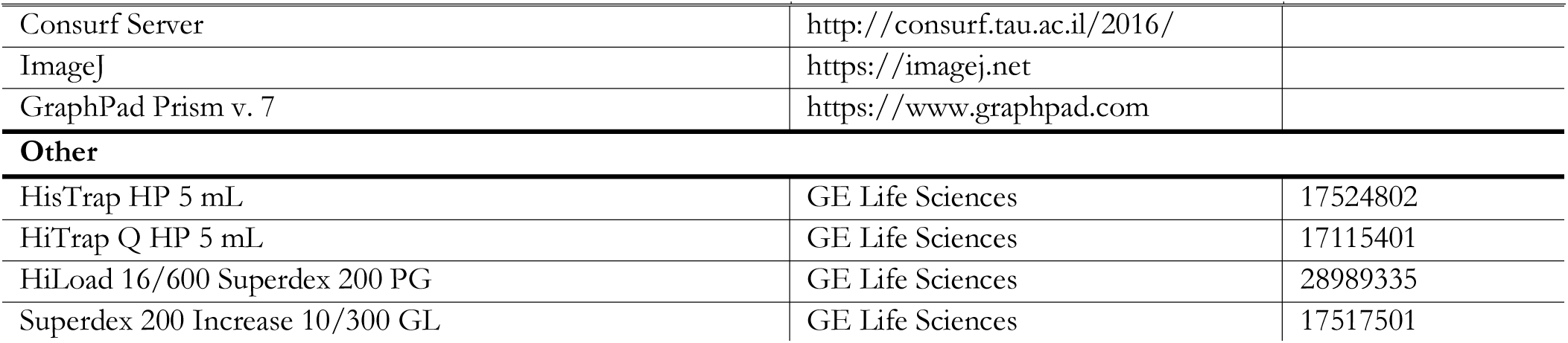
Key Resources

